# Induction of long-term allogeneic cell acceptance and formation of immune privileged tissue in immunocompetent hosts

**DOI:** 10.1101/716571

**Authors:** Jeffrey Harding, Kristina Vintersten-Nagy, Maria Shutova, Huijuan Yang, Jean Kit Tang, Mohammad Massumi, Mohammad Izaidfar, Zohreh Izadifar, Puzheng Zhang, ChengJin Li, Andras Nagy

## Abstract

A vast number of diseases could be treated with therapeutic cells derived from pluripotent stem cells (PSCs). However, cell products that come from non-autologous sources can be immune rejected by the recipient’s immune system. Here, we show that forced expression of eight immunomodulatory transgenes, including *Ccl21, Pdl1, Fasl, Serpinb9, H2-M3, Cd47, Cd200*, and *Mfge8*, allows mouse embryonic stem cells (mESCs) and their derivatives to escape immune rejection in fully immunocompetent, allogeneic recipients. Despite no genetic alterations to major histocompatibility complex (MHC) genes, immune-modified C57BL/6 mESCs could generate long-term, allogeneic tissues in inbred FVB/N, C3H, and BALB/c, as well as outbred CD-1 recipients. Due to the tandem incorporation of our safe-cell suicide system, which allows tight and drug-inducible control over proliferation *in vivo*, these allo-tolerated cells can generate safe and dormant ectopic tissues in the host. We show that these ectopic tissues maintain high expression of all eight immunomodulatory transgenes and are immune-privileged sites that can host and protect unmodified mouse and human cells from rejection in allogeneic and xenogeneic settings, respectively. If translated to human clinical settings, we envision the development of a single pluripotent cell line that can be used to generate allo-tolerated, off-the-shelf cell products to serve all humankind, as well as immune-privileged ectopic tissues to host and immune-protect any kind of therapeutic cell product.

## INTRODUCTION

Immune rejection of allogeneic cells remains a major barrier to the clinical advancement of cell-base therapies. While induced pluripotent stem cells (iPSCs) could be used to generate personalized therapies, routine generation of patient-specific iPSCs in the clinic is unrealistic given the costs and regulatory compliance required. Furthermore, this approach would exclude patients in need of immediate treatment with acute diseases or injuries. Cell banks of PSCs lines selected for the most common human leukocyte antigen (HLA) haplotypes have been suggested as a solution^1^. However, these would fail to serve uncommon haplotypes and still contain differences in minor histocompatibility antigens. While current regiments of immunosuppressive drugs support acceptance of allogeneic cells, systemic immune suppression often leads to life-threatening malignancies and infections.

A potential solution involves the immune modification of PSCs such that their therapeutic derivatives are no longer recognized or attacked by the immune system in allogeneic recipients. A recent report showed that replacement of HLA class I in human PSCs with the inhibitory HLA-E prevented NK and CD8 T-cell responses *in vitro* as well clearance after engraftment into humanized mouse models^2^. Others have combined deletion of MHC class I and II with the forced expression of one or more immunosuppressive transgenes^3, 4^. While promising, MHC-modified or null cells still express minor antigens and other gene polymorphisms that can become targets of rejection and humoral immunity over time^5-7^. Also, cells without MHC/HLA class I or II genes would be unable present viral or bacterial peptides during infection.

We sought to engineer PSCs in a way that accounted for both the major and minor antigens, did not involved deletion of MHC class I/II genes, and which allowed for long-term acceptance in a variety of recipient genetic backgrounds. We were encouraged by prototypical examples of immune escape in nature, such as the devil facial tumour disease (DFTD) type 2, a horizontally-transmitted cancer in Tasmanian Devils^8-10^. Here, we show that the forced expression of eight immunomodulatory transgenes alone in mouse embryonic stem cells (mESCs), without deletion of MHC genes, is sufficient to protect their cell derivatives long-term from rejection in many immunocompetent and allogeneic recipient strains. The use of immunomodulatory factors that interfere with the activity of several immune cell types, including T-cells, NK-cells, antigen presenting cells, and macrophages, was rationalized based on the known complexity of allogeneic immune responses, which may not be reflected in short-term and/or humanized mouse models of allo-transplantation.

Cells that evade immunity system are inherently risky, since the immune system may not be able to clear them should a malignancy arise. Thus, the allo-tolerated ESCs generated here also contain our recently developed safe-cell system^11^. This system consists of a kill-switch genomically-integrated into a gene essential for cell division, providing a unique level of safety for the allo-tolerated cells. It also allows for the generation of a stable, ectopic tissue that can be derived from safe-cell ESCs when the safe-cell system is activated after engraftment *in vivo*. We show that when an ectopic tissue is generated from allo-tolerated mESCs, it becomes a locally-immune-privileged site that can host and protect wild-type allogeneic mouse and xenogeneic human cells from rejection in fully immunocompetent recipient strains.

## RESULTS

### Selection of immunomodulatory genes to disrupt allogeneic immune rejection

To target dendritic cells (Fig. 1a), we selected the gene *Ccl21* encoding a chemokine expressed in the lymph nodes that recruits activated dendritic cells to prime adaptive immunity^12^. Some cancers express CCL21^13^, which may disrupt the ability of local dendritic cells to emigrate out and activate adaptive immunity. To target T-and NK-cells, we selected *Pdl1, Fasl, H2-M3*, ands *Serpinb9* (Fig 1b). PDL1 is a well-described checkpoint inhibitor and can shut down effector T-cells^14^, while FASL can induce T-cell aptopsis^15^. H2-M3 is a non-classical MHC molecule that can inhibit NK cells^16^, while SERPINB9 blocks the apoptosis-inducing Granzyme B released from activated NK and T-cells^17^. To target monocytes and macrophages, we selected *Cd47, Cd200*, and *Mfge8* (Fig 1c). CD47 and CD200 inhibits release of inflammatory cytokines and phagocytosis by macrophages^18, 19^, while MFG-E8 can similarly skew macrophages towards an anti-inflammatory profile during macrophage phagocytosis and contact with dying cells^20^.

**Fig. 1.**
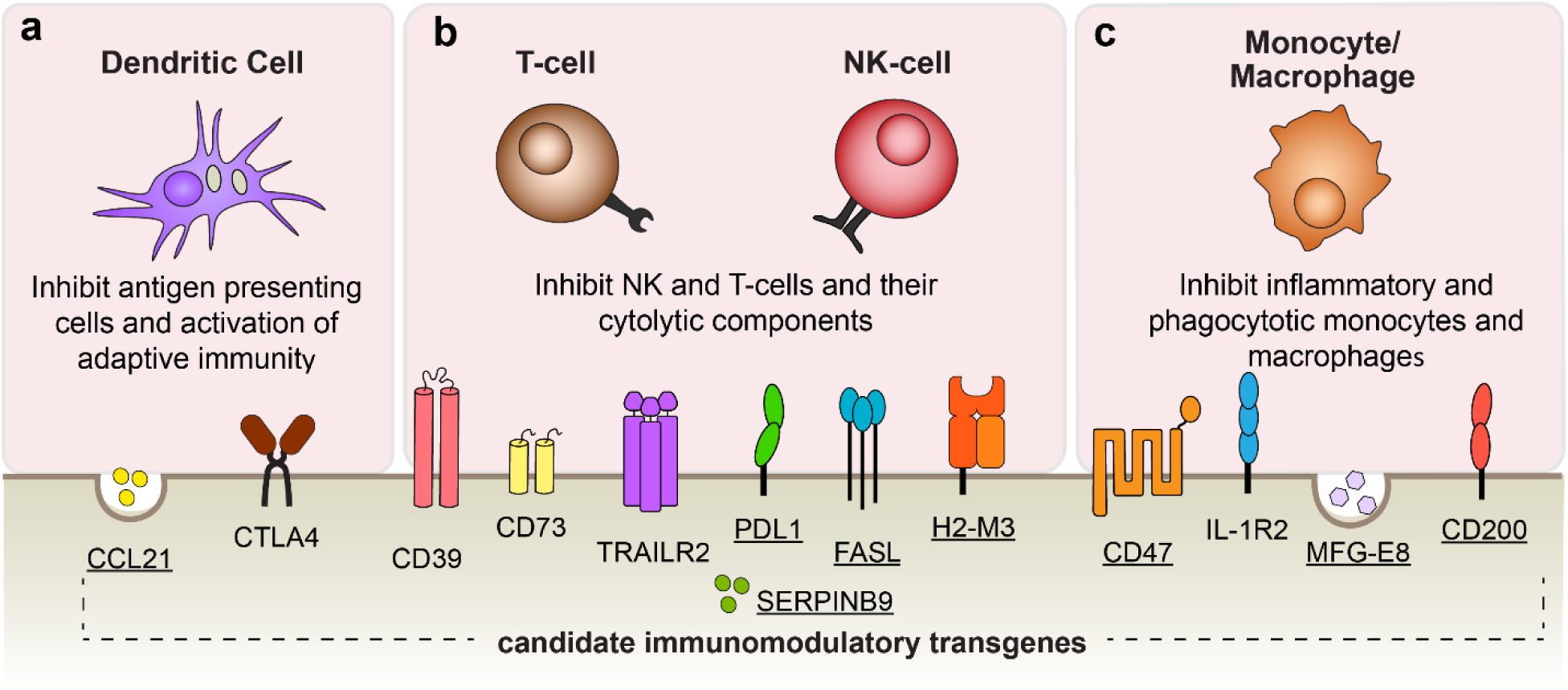
Candidate transgenes and their primary immunomodulatory effect. a. Inhibit the initiation of adaptive immunity and antigen presenting cells, **b**. Inhibit cytotoxic activity of T-cells and NK-cells and their cytolytic components, **c**. Inhibit inflammatory and phagocytotic monocytes and macrophages. Genes from each category (underlined) were selected and introduced into mESCs in order to generate an allo-tolerated line.

### Expression of immunomodulatory transgenes in safe-cell mESCs

The cDNA of the genes encoding these factors were individually cloned into *piggyBac*^21-23^ and *Sleeping Beauty*^*^24, 25^*^ transposon-based expression vectors in which transcription was driven by the strong synthetic CAG promoter^26^ and linked to a drug-selectable (Puromycin or Neomycin) marker or fluorescent (eGFP) reporter by an internal ribosomal entry site (IRES) (Fig. 2a and Supplementary Fig. 1Sa). All eight immunomodulatory transgenes, as well as a transgene encoding enhanced luciferase to allow *in vivo* tracking by Bioluminescent Imaging (BLI), were transfected into mouse safe-cell C57BL/6 mESCs. The safe-cell mESC parental line is a recently-developed clonal derivative of a C57BL/6 mESC line^27^ that contains a thymidine kinase (TK) suicide gene transcriptionally linked to the cell division essential, endogenous *Cdk1* gene^11^. Any dividing cell that co-expresses CDK1 and TK can be killed by activating the TK protein with the pro-drug ganciclovir (GCV).

**Fig. 2.**
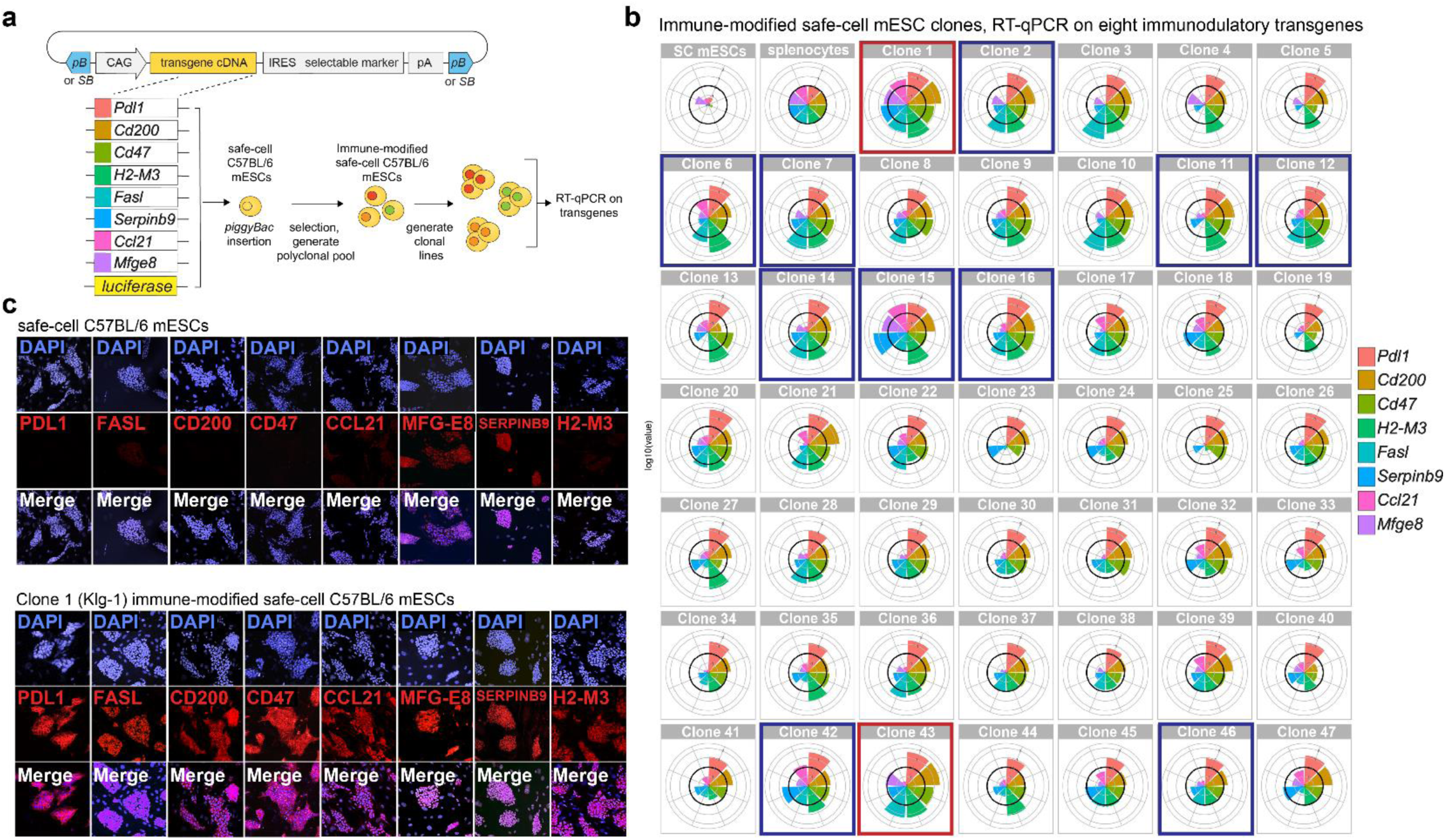
Generating transgenic mESCs expressing eight immunomodulatory genes. Schema of *piggyBa*c (*pB)* and *Sleeping Beauty (SB)*-transposable plasmid used to express immunomodulatory factors in mESCs. cDNA encoding *Ccl21, Pdl1, Fasl, Serpinb9, H2-M3, Cd47, Cd200, and Mfge8* were each cloned into separate *pB* and *SB* transposon vectors and integrated into the genome of safe-cell C57BL/6 mESCs^11^ by transposition in sequential transfection and selection (as described in Supplementary Fig. 1). From the polyclonal transgenic pools, 47 clonal lines were isolated and expanded Expression levels of immunomodulatory transgenes in the clonal lines as measured by RT-qPCR. Each colored wedge shows the value of a specific transgene. All values relative to the expression in isolated splenocytes from C57BL/6 mice (control, thick black circle in each radial graph) activated *ex vivo* with antibodies against CD3εand CD28. Concentric circles; log10 scale. Non-immune modified safe-cell mESCs (parental line) shown in upper left box. Clones outlined in red and blue boxes were later screened *in vivo* for acceptance or rejection in allogeneic hosts (Fig 3). Red boxes; clones accepted in allogeneic hosts (clone 1 and clone 43 renamed Klg-1 and Klg-2, respectively). Blue boxes; clones rejected in allogeneic hosts. **c**. Expression of transgene-encoded factors in Clone 1 (Klg-1) mESCs.

Using the *piggyBac and Sleeping Beauty* transposon system coupled with *in vitro* drug selection and flow cytometry cell sorting, we isolated polyclonal pools of cells with high expression levels of each immunomodulatory transgene (described in Supplementary Fig. 1b). From these polyclonal pools, we then established 47 single cell-derived clonal lines and quantified the expression level of each transgene by qRT-PCR (Fig. 2b). Expression of the protein encoded by each transgene was confirmed with immunohistochemistry on selected clones *in vitro* (Fig. 2c). These factors were undetectable in parental safe-cell mESCs, with the exception of MFG-E8 and SERPINB9, which is consistent with our qRT-PCR results or previously-published results^28, 29^.

### Long-term acceptance of immune-modified C57BL/6 mESCs and their derivatives in allogeneic recipients

From the 47 immune-modified, transgene-containing safe-cell mESC lines, we selected 13 candidate lines with varying overall transgene expression levels, chosen by qualitative assessment of the qPCR radial graphs (Fig. 2b), and used BLI to examine short-term survival in immunocompetent isogenic and allogeneic recipients. As expected, non-immune-modified C57BL/6 (H-2k^b^) safe-cell mESCs survived in isogenic recipients (Fig. 3a, top row, Fig. 3d,e) but were quickly cleared from allogeneic FVB (H-2k^q^), C3H (H-2k^k^), and outbred CD-1 mice within 5-10 days after subcutaneous injection (Fig. 3a). However, some of the generated clones -one hereafter named Klg-1 (shown as “Clone 1” in Fig. 2b) -could still be detected in allogeneic FVB recipients at least 17 days after injection (Fig. 3b). Encouraged by these results, we then tested if Klg-1 mESCs could form teratomas in allogeneic recipients (Fig. 3c). Indeed, Klg-1 mESCs formed well-vascularized teratomas in all allogeneic backgrounds tested, including FVB/N, C3H, BALB/c and CD-1 (Fig. 3d,e). Within two months after transplantation, the mESC-derived teratomas reached a size that required euthanasia of the recipient (Fig. 3d, top graph), which would typically limit the length of such a survival study. So, to test if the Klg-1-derived, allotolerated tissues could survive long-term, we activated the TK-based safe-cell system with the administration of GCV to recipients when the tissues reached an approximate size of 400-500 mm^3^. 14-21 days of treatment with GCV kills all proliferating cells and stabilizes the teratoma tissue indefinitely^11^. We were able to generate long-term and safe-cell-activated Klg-1-derived teratomas in more than 100 allogeneic recipients among all genetic backgrounds tested (Fig. 3d,e). Strikingly, we did not observe a single instance of clearance once these tissues had formed, including mice that were monitored for up to 9 months after transplantation. A complete list of all tested mice and the observation period is shown in Supplementary Table S1. In addition to clone Klg-1, we isolated a second immune-modified clone, termed Klg-2 (shown as “Clone 43” in Fig. 2b), that could also form long-term and GCV-stabilized teratomas in allogeneic recipients, albeit with varying efficiency compared to Klg-1, and which was only tested in FVB and C3H strains (Supplementary Fig. 2).

**Fig. 3.**
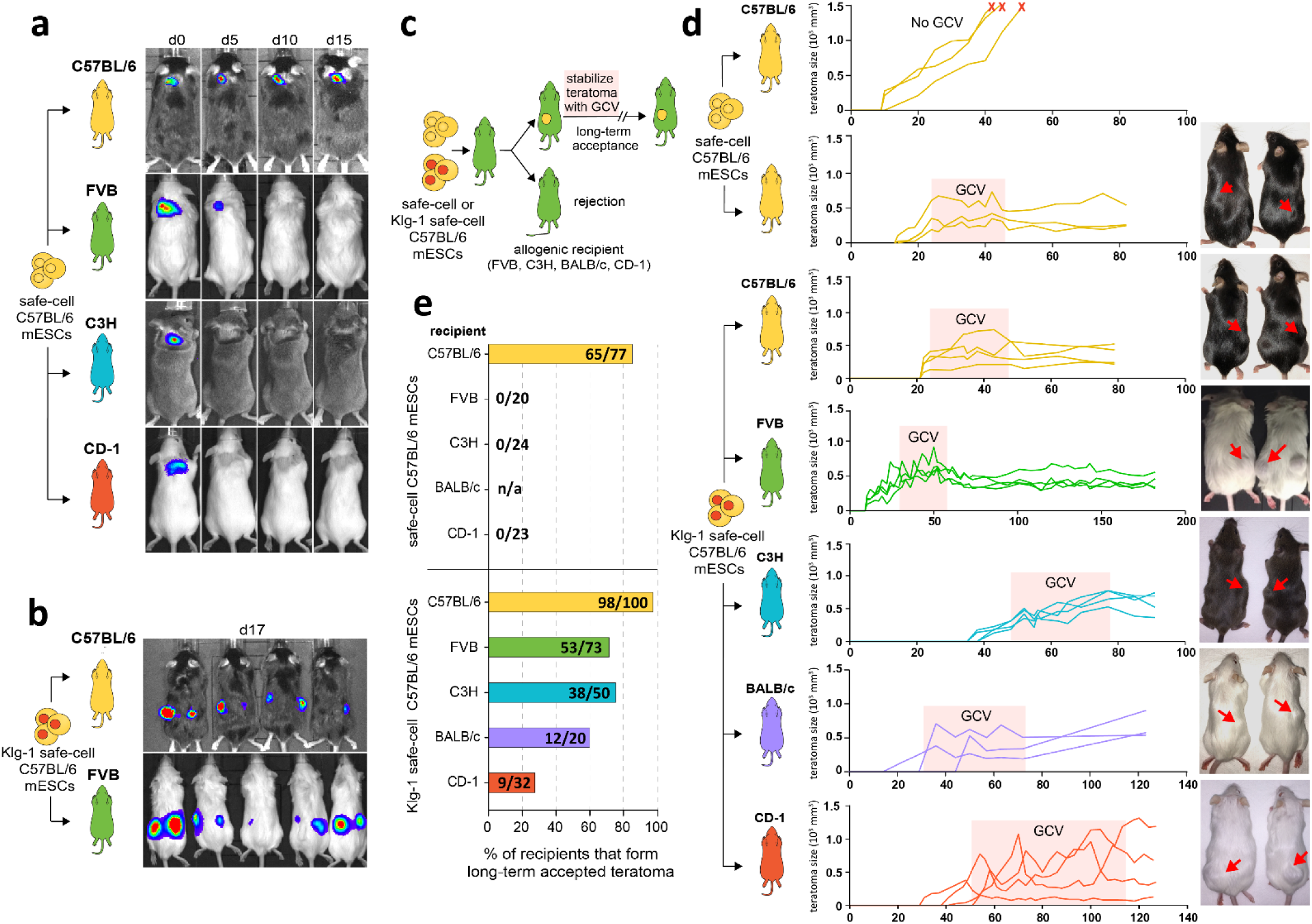
mESCs expressing eight immunomodulatory factors escape immune rejection and survive long-term in multiple MHC-mismatched, immunocompetent strains. **a**. Non-immune modified luciferase+ C57BL/6 safe-cell mESCs were injected subcutaneously into isogenic (C57BL/6) or allogeneic (FVB, C3H, or CD-1) recipients and followed for 15 days with BLI. **b**. Klg-1 mESCs could still be detected in allogeneic (FVB) recipients 17 days after transplantation by BLI. **c-e**. Long-term acceptance of Klg-1 mESCs in isogenic and allogeneic mice. The recipients were monitored for the appearance of teratomas. When the expanding graft reached 500mm^3^, GCV was administered to recipients for several weeks to activate the safe-cell system and ablate dividing cells. Once stabilized, teratomas were monitored for long-term persistence. **c**. schema of long-term acceptance teratoma assay. **d**. Growth and survival curves of untreated and GCV-stabilized teratomas derived from safe-cell and Klg-1 mESCs in isogenic (C57BL/6) and allogeneic (FVB, C3H, BALB/c, and CD-1) recipients. Teratoma growth curves in 3-4 recipients shown per group. Pictures of teratomas (red arrows) shown from two mice per group. **e**. Percent of isogenic and allogeneic recipients that formed long-term and GCV-stabilized teratomas from transplanted safe-cell and Klg-1 safe-cell mESCs. Number in each horizontal bar indicates total number of mice that formed teratomas divided by total number of mice injected. Observation period ranged from 2 to 9 months depending on when recipients were euthanized. No instances of Klg-1 teratoma clearance occurred in any recipient. Data for all mice, including observation period, shown in Supplementary table S1.

Isogenic and allogeneic Klg-1 mESCs teratomas formed cell types of all three embryonic germ layers, including endoderm, mesoderm, and ectoderm (Fig. 4a), indicating that the forced expression of the eight immunomodulatory transgenes did not alter the gross pluripotent developmental potential of the parental cell line. As expected based on their long-term survival, these teratomas were well-vascularized (Fig. 4b). The differentiated cells contained in the GCV-stabilized teratomas expressed high levels of MHC class I molecules, comparable to teratomas derived from non-immune-modified parental safe-cell mESCs in isogenic recipients (Fig. 4c). The robust expression of class I on teratomas demonstrates that the allo-protective effect of the eight immunomodulatory transgenes remains efficient accross MHC differences and can not be attributed to downregulation of MHC.

**Fig. 4.**
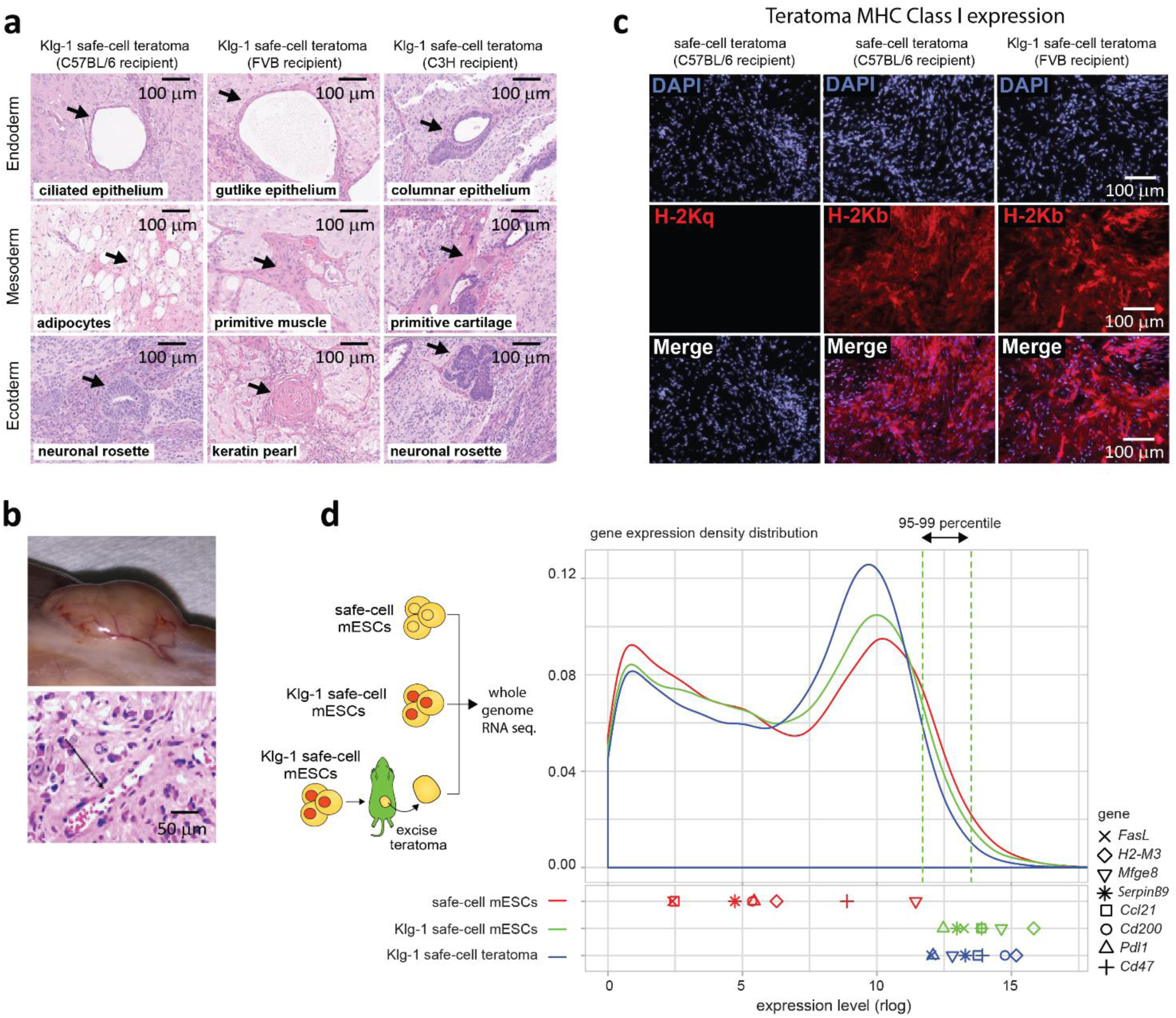
Allo-tolerated Klg-1 mESCs are pluripotent *in vivo*, retain expression of MHC class I and high levels of immunomodulatory transgenes. **a**. histology of long-term Klg-1 derived teratomas in isogenic and allogeneic recipients showing cells derived from all three embryonic germ layers. **b**. vascularization in long-term Klg-1 mESC-derived teratomas formed in allogeneic FVB recipients. **c**. Expression of MHC class I Klg-1 (H-2k^b^) in long-term safe-cell and Klg-1-derived teratomas. MHC Class I H-2Kq serves as negative control for the C57BL/6 background.. **d**. Whole genome RNA sequence gene expression profile from safe-cell (red line and symbols) and Klg-1 mESCs (green line and symbols), as well as cells isolated from long-term Klg-1-derived teratoma in allogeneic FVB recipients (blue line and symbols). Plot shows the proportion of genes (y-axis) at a given expression level (x-axis). Horizontal green dotted lines show 95-99% range.

To quantify and better understand the expression level of each immunomodulatory transgene in reference to the rest of the genome, we then performed next-generation sequencing (NGS) on RNA isolated from *in vitro-*cultured parental safe-cell and Klg-1 mESCs, as well as cells from long-term accepted Klg-1 tissue formed in allogeneic FVB recipients. As expected, the immunomodulatory transgenes were expressed at high levels in Klg-1 mESCs compared to parental safe-cell mESCs, and this high level of expression was preserved in teratomas formed by Klg-1, despite the random cell differentiation that occurs during teratoma development (Fig. 4d). In fact, placed against all the endogenous genes of the genome, the immunomodulatory transgenes in both cultures (Klg-1 mESCs and allogeneic Klg-1 teratomas) were among the highest 5% expressed, with several of the transgenes among the highest 1%. This data fits with the fact that, among the 13 immune-modified mESCs clones tested *in vivo* with a range of overall transgene expression levels, those with the overall highest level of expression (Clone 1 and 43, Fig. 2b) were those that could be allo-tolerated.

We then performed splinkerette PCR on genomic DNA from Klg-1 mESCs using primers against the transposon inverse terminal repeats and sequenced the products by NGS. By comparing the amplified sequence of the products to a reference mouse genome^30, 31^, we identified 56 genomic insertion sites of *piggyBac*-flanked transgenes into Klg-1 mESCs (Supplementary Table S2). Of these insertions, one was in an exon (Hells), one in the 3’UTR (Fasl), and one in the 5’UTR (Anapc11) of a gene. We found that eight intronic insertions and the rest (45) were intergenic.

### Allo-tolerated safe-cell mESCs can generate clinically-relevant cell types *in vitro*

We next examined the *in vitro* properties of Klg-1 mESCs. As expected based on their pluripotency *in vivo*, Klg-1 mESCs had typical ESC colony morphology (Fig. 5a), could form embryoid bodies (Fig. 5b), and expressed the ESC markers OCT4 and SSEA1 (Fig. 5c). In spontaneously-differentiated Klg-1 embryoid bodies, both neuronal and muscle cells could be identified. Using directed protocols, Klg-1 could be differentiated into cardiomyocytes (Fig. 5c, Supplementary Fig. 3, and Supplementary Movie S1,2), CD31+ cells, and Fox2A+Sox17+ cells of the definitive endoderm lineage (Fig. 5c). Together, these data show that forced expression of these eight immunomodulatory factors did not affect Klg-1 mESC pluripotency or the ability to form potentially therapeutic cell types *in vitro*.

**Fig. 5.**
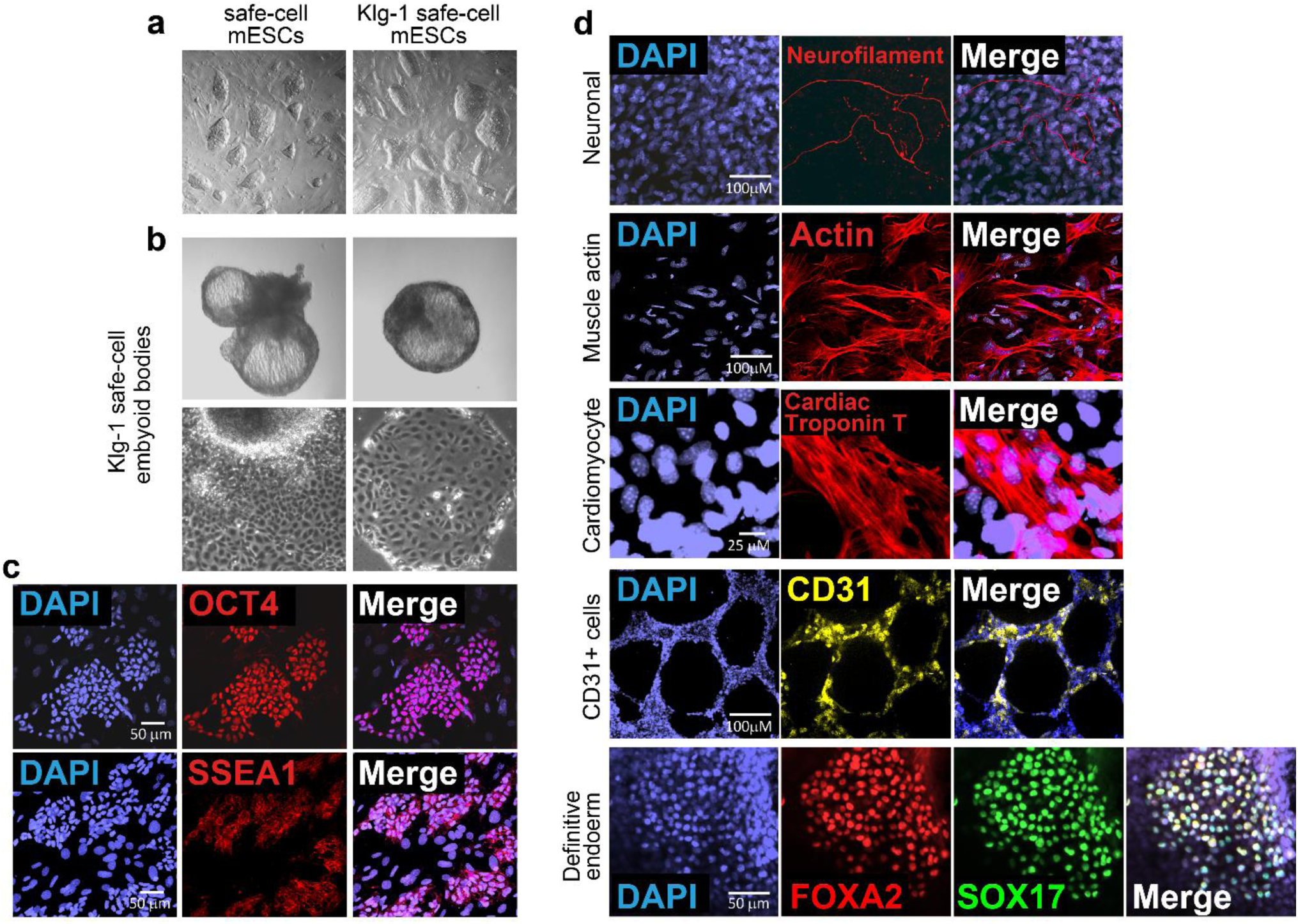
Allo-tolerated Klg-1 mESCs are pluripotent and can generate therapeutic cell types *in vitro*. **a**. Brightfield images of representative safe-cell and Klg-1 mESC colonies *in vitro*. **b**. Embryoid bodies formed from Klg-1 mESCs. **c**. OCT4 and SSEA1 expression in Klg-1 mESCs *in vitro*. **d**. Klg-1 mESCs differentiated into neuronal and muscle cells, as well as cardiomyocytes, CD31+ cells, and definitive endoderm. Functional characterization of cardiomyocytes shown in Supplementary Fig. S4 and Supplementary movie S1,2.

### Ectopic tissues formed by allo-tolerated mESCs serve as an immune-privileged site that can protect wild type allogeneic mouse and xenogeneic human cells

The high level of transgene expression detected at the RNA level in the teratoma (Fig. 4d) was also reflected at their encoded protein levels (Fig. 6a). Based on this observation, we next asked if allogeneic teratomas could host and protect non-immune-modified safe-cells from rejection in allogeneic hosts. First, Klg-1 teratomas were generated in FVB recipients and stabilized by GCV treatment for 40 days (Fig. 6b). Then, allogeneic, non-immune-modified, luciferase+ safe-cell mESCs were injected directly into the teratoma and their presence followed for 40 days by BLI. Injected safe-cell mESCs survived inside the FVB-hosted Klg-1 teratomas and continued to proliferate such that the recipients required an additional round of GCV to stabilize the growth of the injected cells. To show that the survival of allogeneic safe-cell mESCs resulted from the immune-privileged nature of the Klg-1 teratoma, the same safe-cell mESCs were then injected subcutaneously into the contralateral flank of the same FVB recipients. While these subcutaneously-injected mESCs could be detected 5 days later, they were later cleared with no signal detected by 30 days. However, the prior safe-cell mESCs cells injected to the Klg-1 teratoma persisted even while the same safe-cell mESC line was rejected from the contralateral flank.

**Fig. 6.**
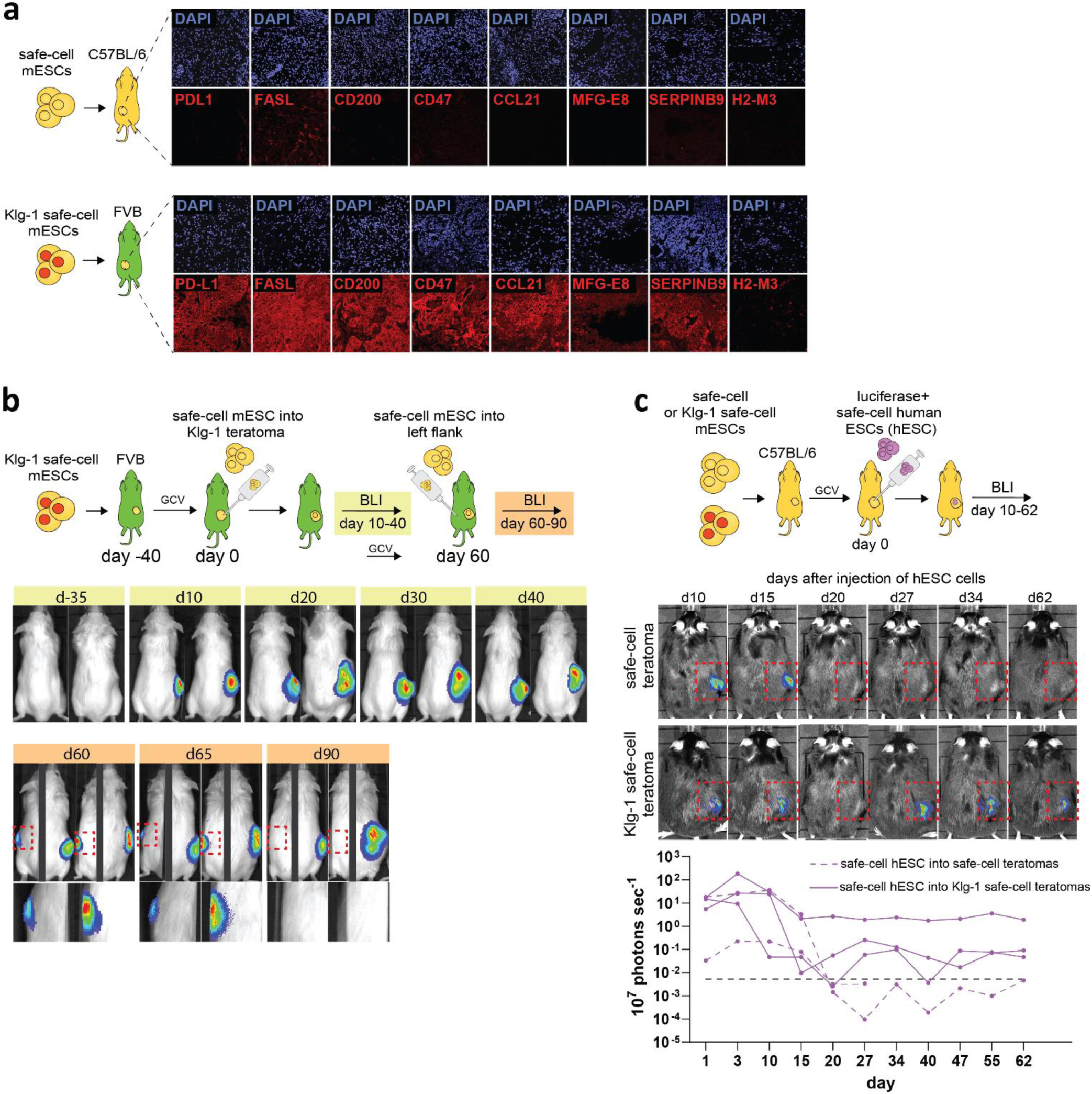
Ectopic subcutaneous tissues formed from allo-tolerated mESCs serve as an immune-privileged site that can protect allogeneic mouse and xenogeneic human cells from rejection. **a**. Antibody staining of immunomodulatory factor in long-term safe-cell and Klg-1-derived teratomas in isogeneic and allogeneic recipients. **b**. Klg-1-derived teratomas protect parental safe-cell mESCs from rejection in allogeneic recipients. Klg-1 teratomas were established and stabilized with GCV over 40 days. 5×10^6^ safe-cell mESCs were then injected into the teratoma and followed with BLI for 40 days (day 10-40). Expansion of safe-cell mESCs within in Klg1 teratoma were halted by GCV treatment. Day-35 image verifies absence of signal detected from Klg-1-derived teratomas using 10 sec exposure time. At day 60, 5×10^6^ safe-cell mESCs were injected subcutaneously into the contralateral (left) flank (red dotted boxes) and followed for 30 days with BLI. BLI images for day 60-90 are composite of two images, each side of the mouse taken at different exposures to account for varying signal intensities (right side; 10 seconds, left side; 1 min). **c**. Klg-1-derived teratomas protect luciferase+ human safe-cell ESCs (hESCs) from rejection in xenogeneic recipients. Stabilized safe-cell or Klg-1 mESC-derived teratomas were established and stabilized with GCV in isogenic C57BL/6 recipients. Then, 10-12 million luciferase+ safe-cell hESCs were injected into the teratoma and followed for 62 days with BLI. Images show one representative mouse, and BLI photon emission of all mice shown graphically (one safe-cell teratoma-injected recipient died from unrelated causes after 27). Red dotted boxes show region containing the teratoma and hESCs. Black dotted line shows detection limit. BLI exposure time;1 min.

We then extended these experiments to a xenogeneic transplant model utilizing our safe-cell human ESC line (hESC^luc+^)^11^. These cells expand and can form GCV-stabilized tissues when injected subcutaneously into immune compromised (Nod-SCID-Gamma, NSG) mice, and can be tracked by BLI (Supplementary Fig. 4). After establishing GCV-stabilized safe-cell or Klg-1 teratomas in isogenic recipients, hESC^luc+^ cells were injected into the teratomas and tracked by BLI for several months. When injected into safe-cell teratomas, hESC^luc+^ cells were cleared within 20 days, but those injected into Klg-1 teratomas persisted long-term without rejection (Fig. 6c). These results show that cells expressing the eight immunomodulatory genes are not only allo-tolerated in a allogeneic setting, but can also form an immune-privileged tissue to protect embedded wild type allogeneic and even xenogeneic cells.

## DISCUSSION

We show here that the forced expression of eight immunomodulatory factors in cells is sufficient to escape immune rejection in allogeneic hosts. Other attempts to engineer allo-tolerated cells have focused on removing MHC genes and/or the introduction of just one or a few factors^2-4, 32^. The emphasis on MHC genes is clearly justified since they are the primary source of allogenic antigens during transplantation and loss of these molecules is often coincident with immune evasion in nature^33^. However, minor antigens, such as those derived from the Y-chromosome or amino acids variations among common proteins can also become targets of immunity, especially during chronic rejection and development of anti-graft antibodies^6, 7, 34^. Removing MHC genes involved in antigen presentation also carry risks, such as the inability to present viral and bacterial peptides should the cells become infected. Escape strategies that rely primarily on deletion of MHC gene also pose the additional risk that they cannot be reversed *in vivo*. The approach here, which relies solely on the use of overexpression of immunomodulatory factors, could be designed into inducible expression systems, such as the TetOFF system, so that transgene expression could be turned off if the transplanted cells become malignant or are no longer needed for a therapeutic effect. This would provide an additional dimension of safety in addition to the use of cell suicide systems.

Our multifactorial transgene approach stems from the premise that immune escape is necessarily difficult due to the highly adaptive and redundant nature of the mammalian immune system, which evolved in an arms race with malignancies, viruses, bacteria, and helminths. In fact, most examples of immune escape in nature require many loss or gain-of-function adaptations. This includes those where MHC expression is retained, including some transmissible cancers^9, 10^, as well as the placenta which expresses HLA-C^35^. The immune modulatory factors used in our study interfere with many key immune pathways, including antigen presentation and the generation of adaptive immunity, NK and T-cell-mediated lysis, and macrophage phagocytosis and general inflammatory responses. These cells and pathways are highly cross-regulated, and targeting them simultaneously may support mutually-reinforcing effects. This is likely an important property of cells that can escape innate, cellular, and humoral immunity over long-term time scales.

The allo-tolerated mESC were generated following modest *in vitro* and *in vivo* screening methods. The necessity of very high expression levels of immunomodulatory transgenes in mESCs and teratomas could serve as a guideline in developing future allo-tolerated cells by similar methods in different species, including human. It emphasizes the importance of strong promoter elements and suitable transgene delivery methods to reduce the amount of *in vitro* and *in vivo* screening needed to identify allo-tolerated lines. In this study, we used a *piggyBac* and *Sleeping Beauty* transposon-mediated transgenesis. Under the proper conditions, we found transposon integration to be so efficient that co-transfection with multiple transgenes resulted in several genomic integration sites of each transgene. These multiple integrations resulted in a high level of expression and mitigated the site-specific silencing and expression variegation that can occur upon differentiation. Random transgene integration does, however, carry the risks of disrupting or activating endogenous genes that could lead to malignancies. It is also true that that the modifications required to generate universally, allo-tolerated cells are inherently dangerous because they can compromise the immune system’s ability to survey and eliminate infected or potentially tumorigenic cells. For both these reasons, it is critical that the cells contain tightly-controlled cell safety systems, such as the one recently published by our group^11^, which was used here. This system consists of an HSV-TK suicide gene integrated into a locus required for cell division and allows reliable removal of dysregulated, dividing cells of graft origin. If needed, both the suicide gene and targeted endogenous gene could be varied to create other types of safe-cell systems, such as one capable of removing all grafted cells regardless of proliferative status, for example. Additionally, our safe-cell system allows for a mathematical quantification of risk^11^, which will be required for patients and clinicians to decide the risk-to-benefit ratio of future cell-based therapies, especially those derived from universal cell products. In addition to allowing engraftment across MHC barriers, the presented approach to achieve allo-tolerance has an additional, unique property. Combined with the safe-cell system, it allowed us to create an immune-privileged tissue that protects not only wild type allogeneic mouse cells but also xenogeneic human cells. After activation of the safe-cell system by GCV treatment of the recipient, the allo-tolerated mESC-derived ectopic tissue is completely stable, dormant, and well-vascularized. Such an “artificial” immune-privileged site could have significant novel therapeutic potential. For example, it could provide a protective site to engraft wild type tissues or organoids that could treat endocrine disease or other factor deficiencies.

The data presented here are the first to demonstrate long-term survival of a solid tissue allograft in fully immune-competent hosts without the need for systemic immunosuppression. These cells’ ability to be allo-tolerated across MHC differences could be the reason behind their status as an immune-privileged tissue. Furthermore, if translated to a human setting our approach satisfies the need for a single pluripotent cell source, which could serve all humankind with safe, off-the-shelf therapeutic cell products and eliminate the need for patients to undergo systemic immune suppression.

## MATERIALS AND METHODS

### Mice

C57BL/6N (Stock No 005304), C3H/HeJ (Stock 000659), FVB/NJ (stock 001800), and BALB/c (Stock 000651) were purchased from Jackson Laboratory. CD1 (ICR) mice were purchased from Charles River. 6-10 week-old mice of each strain or background were used in teratoma assay. Mice were housed in a pathogen-free facility at the Toronto Centre for Phenogenomics (TCP). Mice were housed in micro-isolator cages (Techniplast) with individual ventilation at maximum of 5 mice / cage on a 12-hour light/dark cycle. All mouse procedures were performed in compliance with the Animals for Research Act of Ontario and the Guidelines of the Canadian Council on Animal Care. Animal protocols performed in this study were approved by the TCP Animal Care Committee. The number of animals used in experiments was determined in accordance with similar studies in the field; due to the nature of most experiments, blinding was impossible since the results are visible.

### Construction of *piggyBac* and *Sleeping Beauty* expression vectors

Plasmids containing the cDNA of murine *Pdl1, Fasl, Cd47, H2-M3, Serbinb9, Mfge8, and Ccl21*, were obtained from the Lunenfeld-Tanenbaum Research Institute Open freezer repository. The plasmid containing the cDNA of murine *Cd200* was obtained from GE Dharmacon. The NCIB gene ID, and vendor information, for each gene is listed in Supplementary Table S3. The cDNAs were cloned into *piggyBac (pB)*^21-23^ and *Sleeping Beauty(SB)*^*^24, 25^*^ transposon expression vectors transposon expression vectors} using the Gateway cloning kit (Thermo Fisher 12535029) per manufacturers protocol. Representative plasmid show in Fig. 2a and Supplementary Fig. 1a. Briefly, *Pdl1, Cd47, H2-M3, Mfge8, Ccl21, and Cd200* were amplified by extension PCR using PrimeStar HS mastermix (Takara R040) and gateway compatible attb1/attb2 sites were added to the 5’end and 3’ ends, respectively. All primers used for gateway cloning listed in Supplementary Table S3. The plasmids containing *Ccl21* and *Serpinb9* cDNA contained cDNA-flanking attb1/attb2 sites and did not require extension PCR. All PCR-amplified transgenes were then recombined into the gateway pDONr221 vector (Thermo Fisher 12536017), and the transgene insertion was sequenced in the resulting entry vector using flanking M13 Forward and M13 Reverse, as well as transgene-specific primers listed in Supplementary Table S4. Finally, all transgene-containing entry clones where recombined into the gateway cloning sites contained in *piggyBac* and *Sleeping Beauty* destination vectors to generate the resultant vectors used for transgene expression in cells. At each cloning step, vectors were transfected into chemically-competent DH5alpha cells (Invitrogen) and colonies selected on LB agar plates containing either kanamycin (Sigma K1377) or ampicillin (Sigma 10835242001). Colonies were grown in LB broth (Wisent 809-060-L), and bacterial plasmid DNA was purified using the Presto Mini Bacterial DNA kit (Geneaid PDH300). All sequencing was done by Sanger sequencing at The Centre for Applied Genomics at Hospital of Sick Children (Toronto, Ontario).

### Cell culture

mESCs were cultured in high glucose DMEM (Gibco 11960-044) + 15% fetal bovine serum (FBS) (Wisent 8098-060-L), leukemia-inhibiting factor (LIF) produced at the Lunenfeld-Tanenbaum Research institute (Sinai Health Systems, Toronto ON), 2mM Glutamax (Gibco 35050-061), 0.1 mM nonessential amino acids (Gibco 11140-050), 1mM sodium pyruvate (Gibco 11360-070), 0.1 mM 2-mercaptoethanol (Sigma M6250) and 50ug/mL Penicillin/Streptomycin (Gibco 14140-122). Cells were maintained on a layer of 40k per cm^2^ of mouse embryonic fibroblasts mitotically inactivated by Mitomycin C (Sigma M4287). Cultures were kept in incubators with 5% CO2 at 37°C. Media was changed every day and cells sub-cultured when sub-confluent every 2-3 days. hESCs were cultured on Geltrex-(Gibco A14133-02) coated plates in mTesR (Stem cell Technologies 85851). hESCs were refreshed with mTesR media daily and maintained at 80% confluency.

### Transfection, selection and cloning of mESCs

mESCs were transfected with expression vectors that contained immunomodulatory cDNA using JetPrime transfection kit (Polyplus Transfection 114-01), per manufacturer’s protocol, on adherent culture dishes at approximately 80% confluency. An episomal plasmid encoding hyperactive PBase (hyPBase) (The Sanger Center, pCMV-hyPBase) or sleeping beauty transposase was included when mESCs were transfected with *piggyBac* and *Sleeping Beauty* plasmids. After transfection, transgene-containing clones were generated using combinations of single cell sorting, drug selection with 150 ug/mL G418 (Invitrogen 10131035) or 1ug/mL Puromycin (Gibco A11138-03), bulk sorting using antibody staining against transgene encoded factors, and *in vitro* culture plating at clonal densities (200 cells per 100mm dish). The transfection sequence, sorting, and drug selection that led to transgene-containing clones is described in Supplementary Fig. S1. For the drug selection steps, G418 or Puromycin was added to the selection media 24 hours after transfection and maintained until the appearance of antibiotic-resistant cells. All final transgene-containing clones were grown in triplicate 96-well plates for cryopreservation and DNA/RNA extraction.

### qRT-PCR of immunodulatory transgenes

RNA was isolated from clonal cultures by removing the media and adding Trizol (Invitrogen 15593-031) for 5 minutes at room temperatures, followed by 150uL Chloroform (Sigma). After briefly vortexting and centrifugation at high speed, the nucleic acid phase was collected, precipitated with 100% iso-propanol, washed with 70% ethanol, and eluted in RNAse free water. cDNA was then made from isolated RNA with the QuantiTect Reverse Transcription Kit (Qiagen 205310). qPCR was performed using the SensiFast SYBR No-Rox Kit (Frogga Bio BIO-98020) per manufacturer’s protocol on a Bio-Rad C1000 CFX384. The forward and reverse primers used to detect inserted transgenes designed to span only exons to avoid detection of endogenous genes. Transgene expression levels were compared to levels of activated splenocytes, and everything normalized to primers against Eef2. All primer sequences listed in Supplementary Table S5.

### Activation of splenocytes

The spleen from a C57BL/6N mouse was manually homogenized in a 70um cell strainer (Falcon 352350) in DMEM (Gibco 11960-044). The single cell suspension was washed with DMEM, lysed with Red Blood Cell lysis buffer (Sigma R7757-100), and washed again with DMEM. Isolated splenocytes were plated onto adherent culture dishes and activated with anti-mouse antibodies against CD3ε(1:1000 dilution, BD Biosciences 553058) and CD28 (1:1000 dilution, BD Biosciences 553295), as well as Golgi Stop (1:1000 dilution, BD Biosciences 554725) for 8 hours in high glucose DMEM (Invitrogen) + 15% fetal bovine serum (FBS) (Gibco 12483-020), 2mM Glutamax (Gibco 35050-061), 0.1 mM nonessential amino acids (Gibco 11140-050), 1mM sodium pyruvate (Gibco 11360-0700) mM 2-mercaptoethanol (Sigma M6250) and 50ug/mL Penicillin/Streptomycin (Gibco 15140-122). RNA was then isolated from activated cultures and used as control in RT-qPCR analysis of transgene-containing mESC clones.

### Whole genome RNAseq

Teratomas were excised, frozen on dry ice, and then homogenized with a TissueLyser II (Qiagen 85300) per manufactuer’s protocol. RNA was then collected from isolated teratoma cells, as well as mESCs grown in-vitro, using the Qiagen RNeasy kit (Qiagen 74104). cDNA was made from isolated RNA with the QuantiTect Reverse Transcription Kit (Qiagen 205310). RNA-Seq was performed using an Illumina HiSeq 2500 (The Center For Applied Genomics, Hospital for Sick Children, Toronto ON). Trimmomatic^36^ was used to trim residual Illumina adapters from reads, and FastQC tool (https://www.bioinformatics.babraham.ac.uk/projects/fastqc/) for quality controls. Trimmed reads were pseudo-aligned with kallisto^37^ and further processed using DESeq2^38^ in R (http://www.r-project.org/). The DESeq2 tool was used to produce rlog values.

### Splinkerette PCR and analysis

Splinkerette PCR (spPCR) was done on genomic DNA from Klg-1 mESCs using Primestar HS mastermix (Takara R040) and primers specific for the inverse terminal repeats of the *piggyBac* sequences flanking the immunomodulatory transgenes. HMSpAa (top strand primer) CGAAGAGTAACCGTTGCTAGGAGAGACCGTGGCTGAATGAGACTGGTGTCGACACTAGTGG; HMSpBb (lower strand primer/hairpin) gatcCCACTAGTGTCGACACCAGTCTCTAATTTTTTTTTTCAAAAAAA Primary PCR primers: PB-L-Sp1 (PB 5’TR sequence) GCGTGCTTGTCAATGCGGTAAGTGTCACTG; PB-R-Sp1 (PB 3’TR sequence) CCTCGATATACAGACCGATAAAACACATGC. Secondary nested PCR primers: PB-L-Sp2 (PB 5’TR sequence) ACGCATGCATTCTTGAAATATTGCTCTCTC; PB-R-Sp2 (PB 3’TR sequence) ACGCATGATTATCTTTAACGTACGTCACAA. Amplified DNA was gel-extracted with the Monarch Gel Extraction Kit (NEB T1020L) and sequenced by next generation sequencing (NGS). Residual Illumina primers, as well as *piggyBac* and spPCR primers were trimmed from reads using Trimmomatic^36^. Trimmed paired reads were then aligned with the end-to-end mode of STAR tool^39^ and visualized in IGV^40, 41^. High quality reads were then filtered for only those beginning or ending with TTAA (the sequences at which *piggyBac* inserts into the genome^23^) and overlapping with the nearest read by a TTAA sequence. 93 candidate regions of insertion were identified, and then filtered to sort on only those where paired reads were present on both sides of the TTAA region and without a TTAA in the middle of the read, ultimately resulting in 56 high quality candidates, listed in Supplementary Table S2.

### Bioluminescent Imaging (BLI)

Mice were injected intraperitoneally with 100ul of 15mg/mL solution of Luciferin (Xenolight D-Luciferin, Perkin Elmer 122799). 15 minutes following injection, mice were anaesthetized with gaseous Isoflurane and imaged with the IVIS Lumina imager (Perkin Elmer) at exposures between 10 seconds and 3 minutes. Binning was set to small and F/stop to 1. Images we acquired using Living Images Software (Perkin Elmer). Short exposure times of 10 seconds were used to identify safe-cell mESCs (which are luciferase^high^) injected into Klg-1-derived teratomas (which come from luciferase^low^ Klg-1 mESCs).

### *In vivo* mESCs injections and teratoma assay

Matrigel matrix High concentration (Corning 356234) was diluted 1:1 in ice-cold DMEM and kept on ice. mESCs were treated with Trypsin-EDTA at 37°C, resuspended in ESC media, centrifuged, and washed in DMEM. Then, 5×10^6^ mESCs were diluted in 100uL of ice-cold Matrigel:DMEM mixture and injected subcutaneously at the back of the neck (for BLI imaging), or each dorsal flank (two separate injections) for teratoma assays (5×10^6^mESCs in 100uL at each flank). Mice were anaesthetized during injections with isofluorane. Palpable teratomas developed 10-40 days after injection, depending on the strain. Teratoma size was measured with calipers and the volume calculated using the formula V = (LxWxH)π/6. At endpoint, mice were euthanized and the teratomas excised for immunohistochemistry and histology.

### Stabilization of teratomas with Ganciclovir (GCV)

All mESCs used in these studies contain the safe-cell system which contains a thymidine kinase suicide gene targeted to the CDK1 locus^11^. Therefore, for long-term acceptance studies, mice with mESCs-derived teratomas were given Ganciclovir (Cytovene) once teratomas reached an approximate volume of 400-500mm^3^ for a period of several weeks, depending on the strain. Any mice in which teratoma growth resumed at a later date after GCV withdrawal was given GCV again for several days. Ganciclovir administration was given as daily intraperitoneal injections at a dose of 50mg/kg in 100uL PBS.

### Flow cytometry and cell sorting

Single cells suspensions of transfected mESCs were collected at 80% confluency after trypsinization, washed with FACS buffer (PBS + 1% BSA and 0.5% EDTA), and stained with primary antibodies against mouse PD-L1 (Novus NBP1-43262), CD47 (BD Biosciences 740055), CD200 (BD Biosciences 565544), and H2-M3 (BD Biosciences 551769).

All antibody cocktails contained 1:100 dilution of anti-mouse Fc Receptor block CD16/CD32 (BD Biosciences 553141). Primary antibodies against H2-M3 was detected with secondary antibody conjugated to Alexa568 (Thermo Fisher A21112) by staining in PBS + blocking buffer at room temperature. Cells were then washed in FACS buffer, sorted using an LSRFortessa (BD Biocsiences) in FACs buffer + 25mM HEPES buffer (Sigma H3537)) and collected in mESC cell media. DAPI (ThermoFisher 62248) was added to all samples prior to sorting to collect only live cells, and doublets excluded by forward / side scatter. Wild-type mESCs stained with the same antibodies and single-color samples were used as negative controls and for instrument compensation. Flow cytometry was performed at the Lunenfeld-Tanenbaum Research Institute flow cytometry facility and the data analyzed using FlowJo v10 (TreeStar).

### Fluorescent Immunohistochemistry

mESCs were grown on mitotically-inactivated MEFs plated onto coverslips coated with gelatin (STEMCELL Technologies 07903). Coverslips were fixed with 4% PFA (Sigma), washed with PBS, and incubated with 100mM glycine (Sigma) at 4 ^o^C in PBS with 0.02% Sodium Azide (Sigma 71289). Coverslips were then blocked with PBS + 10% donkey serum, 0.01% triton, and 1:100 anti-mouse CD16/CD32 (BD Biosciences 553141) and stained with primary antibodies in PBS + 10% donkey serum. For immunohistochemistry of teratoma tissue sections, teratomas were fixed in 4% PFA at room temperature for 24-48 hours, dehydrated in 25% sucrose in PBS, embedded in OCT compound, and cryo-sectioned. Tissue sections were fixed in cold acetone for 10 minutes, washed with PBS, blocked in PBS + 3% BSA, 10% goat serum and 1:100 anti-mouse CD16/CD32, and then stained with primary antibodies overnight at 4°C in blocking buffer. For both mESCs on coverslips and teratoma tissue sections, primary antibodies were used against mouse PD-L1 (Novus NBP1-43262), CD47 (BD Biosciences 740055), CD200 (BD Biosciences 565544), H2-M3 (BD Biosciences 551769), FASL (Novus NBP1-97519), SERPINB9 (Hycultbiotech HP8035), CCL21 (R&D Systems AF457), and MFGE8 (Biolegend 518603). Additional primary antibodies used on *in vitro*-cultured cells include anti-mouse CD31 (Biolegend 102417), Neurofilament (Fisher Scientific MS359R7), SSEA1 (Sigma MAB4301), Smooth Muscle Actin (Abcam ab5694), FOXA2 (Abcam ab108422), and SOX17 (R&D Systems AF1924), and anti-cardiac Troponin-T (cTNT)(abcam, ab8295). Other primary antibodies used on tissue sections included anti mouse H-2Kb (Biolegend 116511), and H-2Kq (Biolegend 115106). Primary antibodies against H2-M3, FASL, SERPINB9, CCl21, MFGE8, Neurofilament, OCT4, SSEA1, Smooth Muscle Actin, FOXA2, and SOX17 were detected with secondary antibodies conjugated to Dylight 488 (Biolegend 405503), DyLight 549 (Novus NBP1-72975), or Alexa647 (Life Technologies 21244) by staining in PBS + blocking buffer at room temperature. mESCs on coverslips and tissue sections were stained with DAPI (ThermoFisher 62248) and mounted with Immunomount (Thermo Fisher 9990402).

### Histology

Paraffin embedding, sectioning and H&E staining was carried out by staff at the Toronto Centre for Phenogenomics Histology laboratory.

### Generation of Embryoid Bodies and spontaneous differentiation

To generate embryoid bodies (EBs) mESCs were collected and plated in suspension on non-adherent tissue cultures plates (Corning, ULC 3471) in mESC media without LIF and FBS (Wisent) concentration reduced to 10%. To induce spontaneous differentiation, EBs were then collected, and plated onto adherent tissue culture plates that were coated with Gelatin (STEMCELL 07903). The presence of cardiomyocytes, as well as cells containing neurofilament and muscle actin were then confirmed within the cells that proliferated and differentiated out from attached EBs.

### Directed differentiation of mESCs into Cardiomyocytes

mESCs were maintained in serum-free (SF) mouse ESC culture medium on irradiated feeders. On day 0, mESCs were dissociated with TrypLE (Invitrogen) and aggregated in SF medium with LIF. After 48 hours, the embryoid bodies (EBs) were dissociated with TrypLE and cultured on matrigel-coated 24-well cell culture plate in SF medium with LIF for 8 h followed by media replacement with SF differentiation medium without LIF for 24 hours. On day 3, 5 ng/mL VEGF (PeptroTech 450-32), 0.25 ng/mL BMP4 (PeptroTech 315-27) and 5 ng/mL Activin A (R&D Systems 338-AC-050/CF) were added to the SF differentiation medium. On day 4, the differentiation medium was replaced by StemPro 34 medium (Invitrogen) 5 ng/mL VEGF (PeptroTech 450-32), 10 ng/mL bFGF (PeptroTech 100-18B), and 50 ng/mL FGF10 (PeptroTech 100-26). The media was changed every 60 hours. On day 15, the contractile mESC-derived cardiomyocytes were examined for contractility and calcium transients. A custom MatLab code was developed to analyze the images of the contractile cardiomyocytes. Immunohistochemistry was performed to stain cells with the primary anti-cardiac Troponin-T antibody (abcam, ab8295) and detected with secondary fluorophore-conjugated antibody against Goat IgG (abcam ab205719).

### Differentiation of mESCs into definitive endoderm

mESCs were aggregated in suspension in mouse mESC cell media supplemented with 3% Knockout Serum Replacement (SR), without LIF to form EBs. The next day (day 1), 100ng/mL Activin A (R&D Systems 338-AC-050/CF) and 75 ng/mL Wnt3a (PeproTech 7315-200) were added to the EB culture media. On day 3, the EB media was replaced with mESC (without LIF) + 3 % SR and 100ng/mL Activin A (100 ng/ml). On Day 5, cells were stained with primary antibodies against SOX17 (R&D Systems AF1924) and FOXA2 (Abcam 108422) and detected with secondary fluorophore-conjugated antibodies against Goat IgG (Thermo Fisher 11055) and Rabbit IgG (Thermo Fisher 31537), respectively.

### Differentiation of mESCs into CD31+ cells

mESCs were disassociated with Accutase (STEMCELL Technologies 07920) and washed in DMEM + 10% FBS (Wisent). Then, cells were plated at 1 million cells per well in 2mL mESC media (without LIF) + 25 μg/L Activin A (R&D Systems 338-AC-050/CF), 5 μg/L BMP4 (PeptroTech 315-27), 1 μM CHIR99021 (R&D Systems 4423/10), and 10 µM Rock inhibitor(STEMCELL Technologies Y-27632) onto 6-well plates coated with 1mL/well of 1µg/cm^2^ Vitronectin XF (STEMCELL Technologies 07180) for 1 hour at room temperature. The media was then replaced each day for the next two days (days 1, 2) with 4mL of mESC media (without LIF) + 10% FBS, 25 μg/L ActivinA, 5 μg/L BMP4, and 1 μM CHIR99021. The media was then replaced daily for the next 4 days (day 3, 4, 5, 6) with 4mL mESC media (without LIF) + 10% FBS, 50 μg/L BMP4, 50 μg/L VEGF-A (PeptroTech 450-32), and 5 μM SB431542 (Sigma S4317). The next day (day 7), cells were disassociated with Accutase, and CD31+ cells were isolated using flow cytometry sorting after staining cells with anti-mouse CD31 antibody (Biolegend 102417) and plated onto Vitronectin-coated plates (1mL/well at 1µg/cm^2^) in mESC media (without LIF) + 10% FBS and 50 μg/L VEGF-A. This media was replaced every other day and the cells passaged when confluent at 1:3 until maturation into endothelial cells.

## Acknowledgements

We thank Drs Claudio Monetti and Qin Liang for providing the safe-cell mESCs, Annie Bang for support with flow analysis, and Ellen Langlille for contributed artwork. We would also like to acknowledge the staff at the TCP animal facility, in particular Christina Dalrymple, for their support. Last but not least, we owe our respect to the mice that have contributed to these studies.

## Author contributions

J.H. wrote the manuscript, contributed to conceptual and experimental design, generated data, and analyzed data. K.N. wrote the manuscript, contributed to conceptual and experimental design, generated data, and analyzed data. H.Y. generated and analyzed data. M.S. analyzed NGS and qRT-PCR data. Z.I., M.M., C.L., and P.Z. generated data. A.N. contributed to conceptual and experimental design, and wrote the manuscript.

## LIST OF SUPPLEMENTARY FIGURES, TABLES, MOVIES

**Supplementary Fig. S1**. Engineering immunomodulatory transgenes into mESCs

**Supplementary Fig. S2**. Immune-modified clone Klg-2 accepted in MHC-mismatched, immunocompetent strains.

**Supplementary Fig. S3**. Characterization of cardiomyocytes differentiated from Klg-1 mESCs

**Supplementary Fig. S4**. Safe-cell hESCs-derived teratomas in NSG mice.

**Supplementary Table S1**. Klg-1 and Klg-2 teratomas in recipient mice

**Supplementary Table S2**. Number and position of *piggyBac* insertions,

**Supplementary Table S3**. Attb1-forward and attb2-reverse primers used for gateway cloning.

**Supplementary Table S4**. Primers used to verify transgene sequence after cloning

**Supplementary Table S5**. RT-qPCR forward and reverse primers

**Supplementary Video S1**. Cardiomyocytes derived from Klg-1 mESCs, contractility

**Supplementary Video S1**. Cardiomyocytes derived from Klg-1 mESCs, calcium flux

---

**Supplementary Fig. S1.**
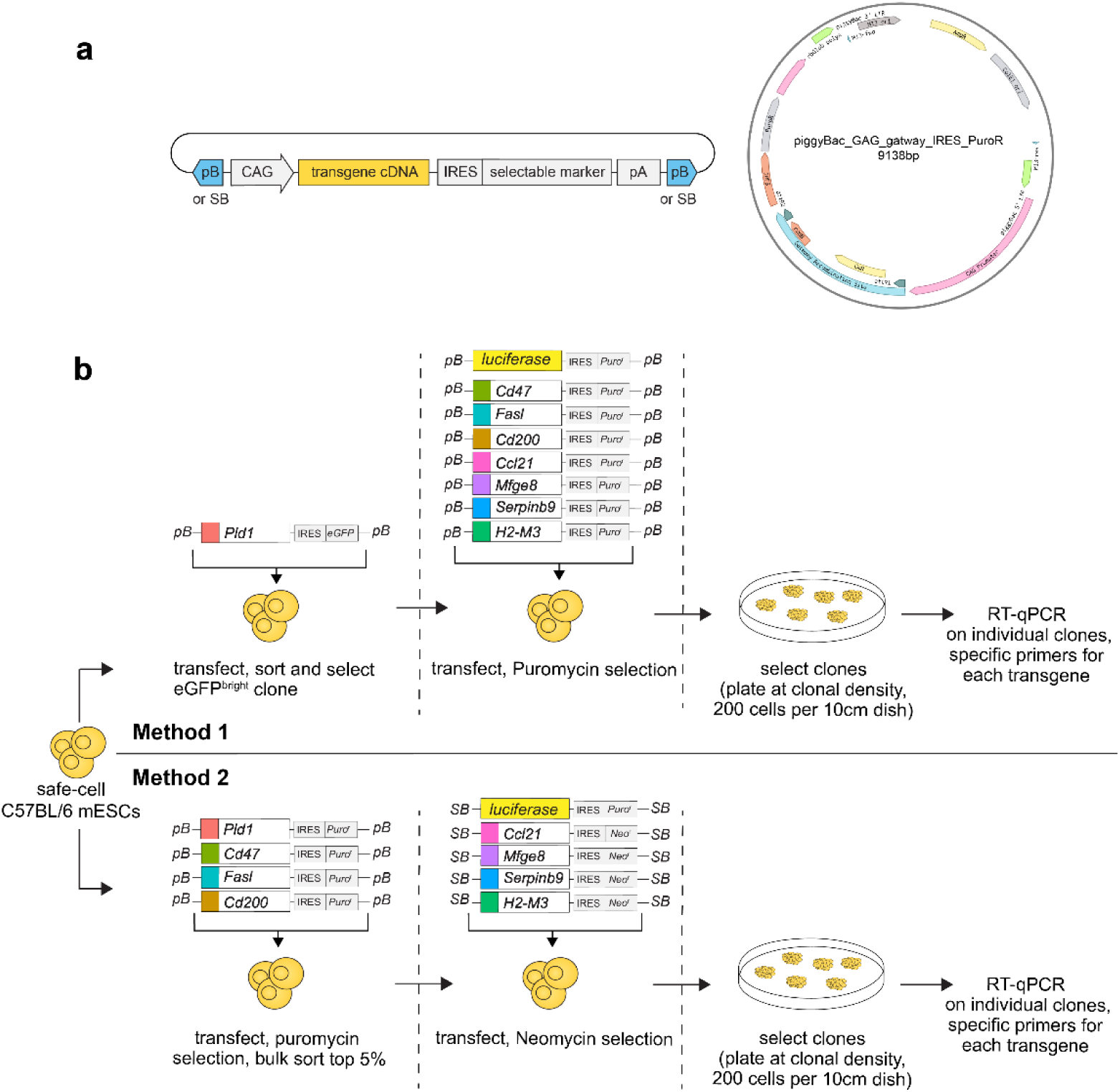
Engineering immunomodulatory transgenes into mESCs. **a**. Map of representative *piggyBac (pB)* and *Sleeping Beauty (SB)* plasmid used to express eight immunomodulatory transgenes (*Ccl21, Pdl1, Fasl, Serpinb9, H2-M3, Cd47, Cd200*, and *Mfge8)* and luciferase in safe-cell C57BL/6 mESCs. Transgene expression driven by a CAG promoter and linked by an IRES to an eGFP fluorophore, or *Puromycin*^*r*^ or *Neomycin*^*r*^ genes. Expression cassette flanked by long-terminal repeat *piggyBac* or *Sleeping Beauty* sequences. **b**. Two sequential methods were used to generate safe-cell mESCs with expression of all eight immunomodulatory transgenes: Method 1) safe-cell C57BL/6 mESCs transfected with eGFP-linked *Pdl1*, eGFP^high^ clone was isolated, followed by simultaneous transfection of *Ccl21, Fasl, Serpinb9, H2-M3, Cd47, Cd200, Mfge8* and *luciferase* (each containing *Puromycin*^*r*^) and drug selection with Puromycin. Method 2) safe-cell C57BL/6 mESCs simultaneously transfected with *Pdl1, Fasl, Cd47, and Cd200*, each with IRES-linked *Puromycin*^*r*^, followed by selection with Puromycin and flow cytometry sorting for the highest 5% expression of each factor using factor-specific antibodies. Bulk sorted cells then transfected with *Ccl21, Mfge8, SerpbinB9*, and *H2-M3* containing *Neomycin*^*r*^, followed by selection with Neomycin. Both methods resulted in a polyclonal pool subcloned by limited dilution plating, Clones were analyzed by RT-qPCR for transgene expression levels (Fig. 2b in main text).

**Supplementary Fig. S2.**
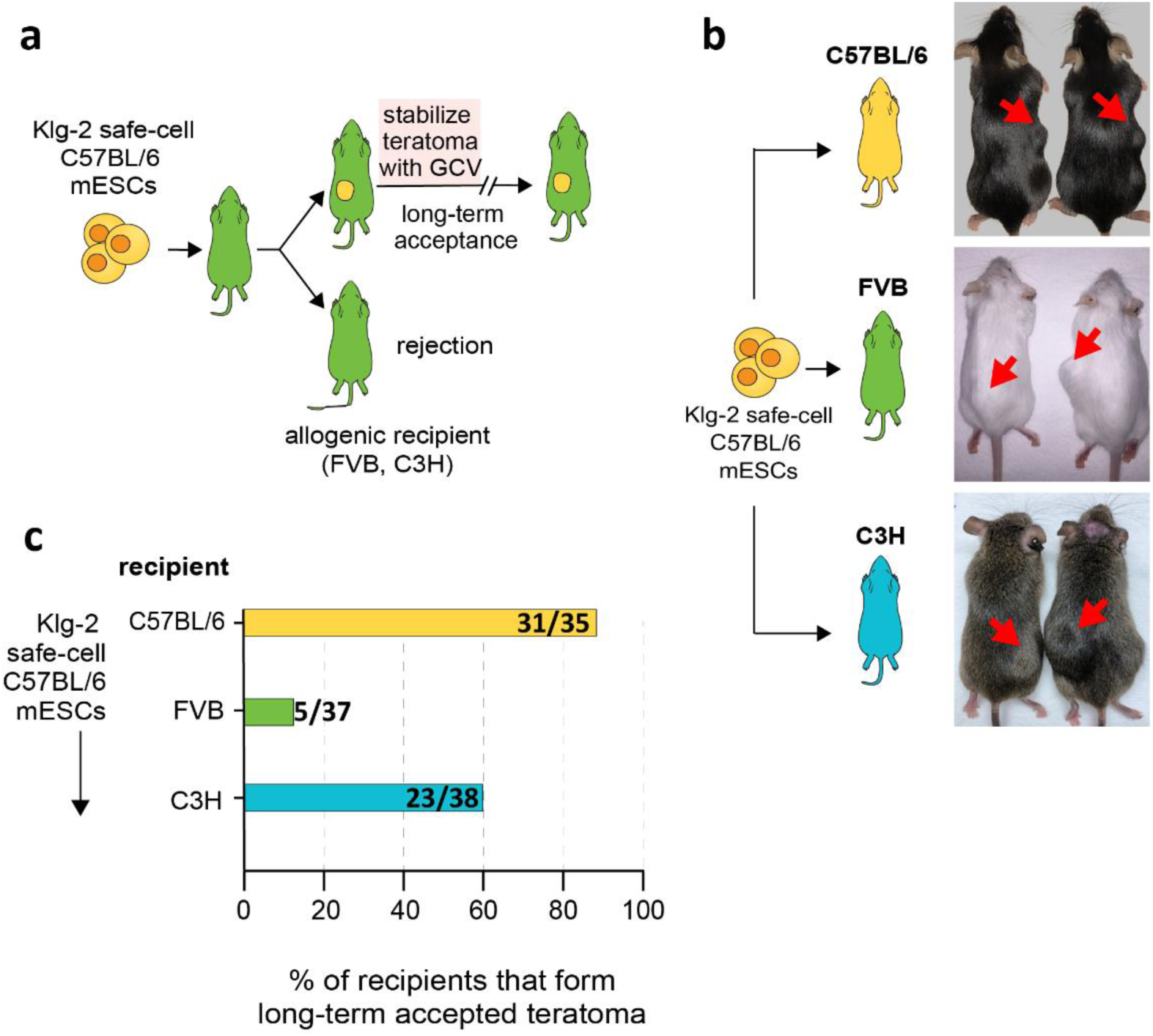
Immune-modified clone Klg-2 accepted in MHC-mismatched, immunocompetent strains. Klg-2 mESCs (renamed from Clone 43 in Fig 2b. of main text, outlined in red) was tested for survival and long-term persistence in a limited number of allogeneic recipient backgrounds. **a**. schema of long-term acceptance teratoma assay. **b**. Pictures of teratomas (red arrows) shown in two mice per group that formed long-term teratomas. **c**. Percent of isogenic and allogeneic recipients that formed long-term, GCV-stabilized teratomas from transplanted Klg-1 mESCs. Number in horizontal bar indicates total number of mice that formed teratomas divided by total number of mice injected. Observation period ranged from 2 to 6 months. No instances of Klg-2 teratoma clearance occurred in any recipient. Data for all mice, including observation period, shown in Supplementary table S1.

**Supplementary Fig. S3.**
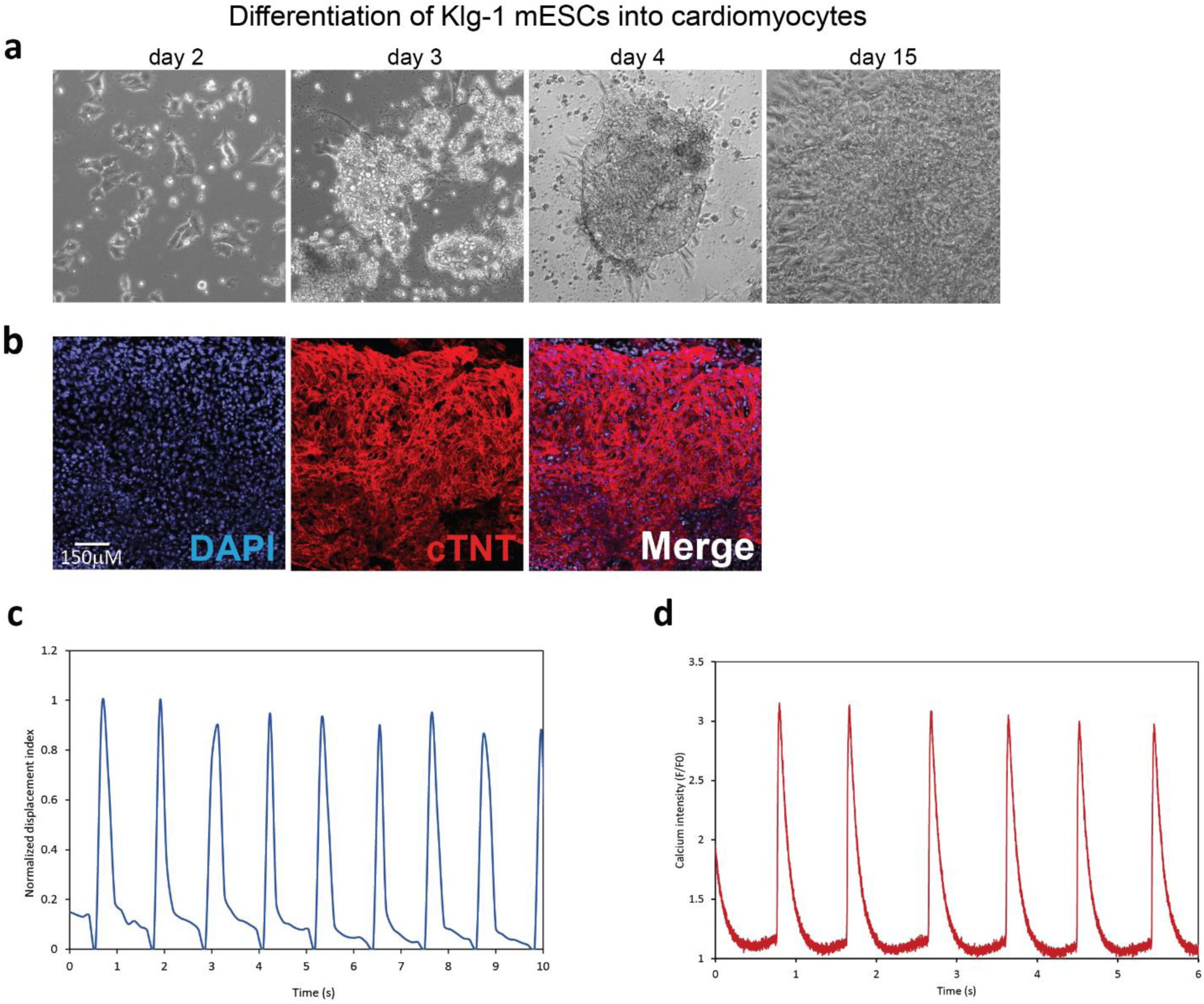
Characterization of cardiomyocytes differentiated from Klg-1 mESCs. **a**. Representative brightfield micrographs of Klg-1 mESCs during directed differentiation into cardiomyocytes. Day 2, adhesion of mESCs following embryoid body dissociation and cell seeding on matrigel-coated cell culture plates. Day 3, proliferation and cluster formation for cardiac mesoderm induction. Day 4, early cardiac differentiation. Day 15, contractile cardiomyocytes. **b**. IHC showing expression of cardiac Troponin T (cTNT) after 15 days of differentiation. **c**. Cardiomyocyte cyclic contraction 15 days after differentiation, measured as a function of normalized cell displacement. **d**. Cardiomyocyte calcium flux after loading with Fluo-4 AM.

**Supplementary Fig. S4.**
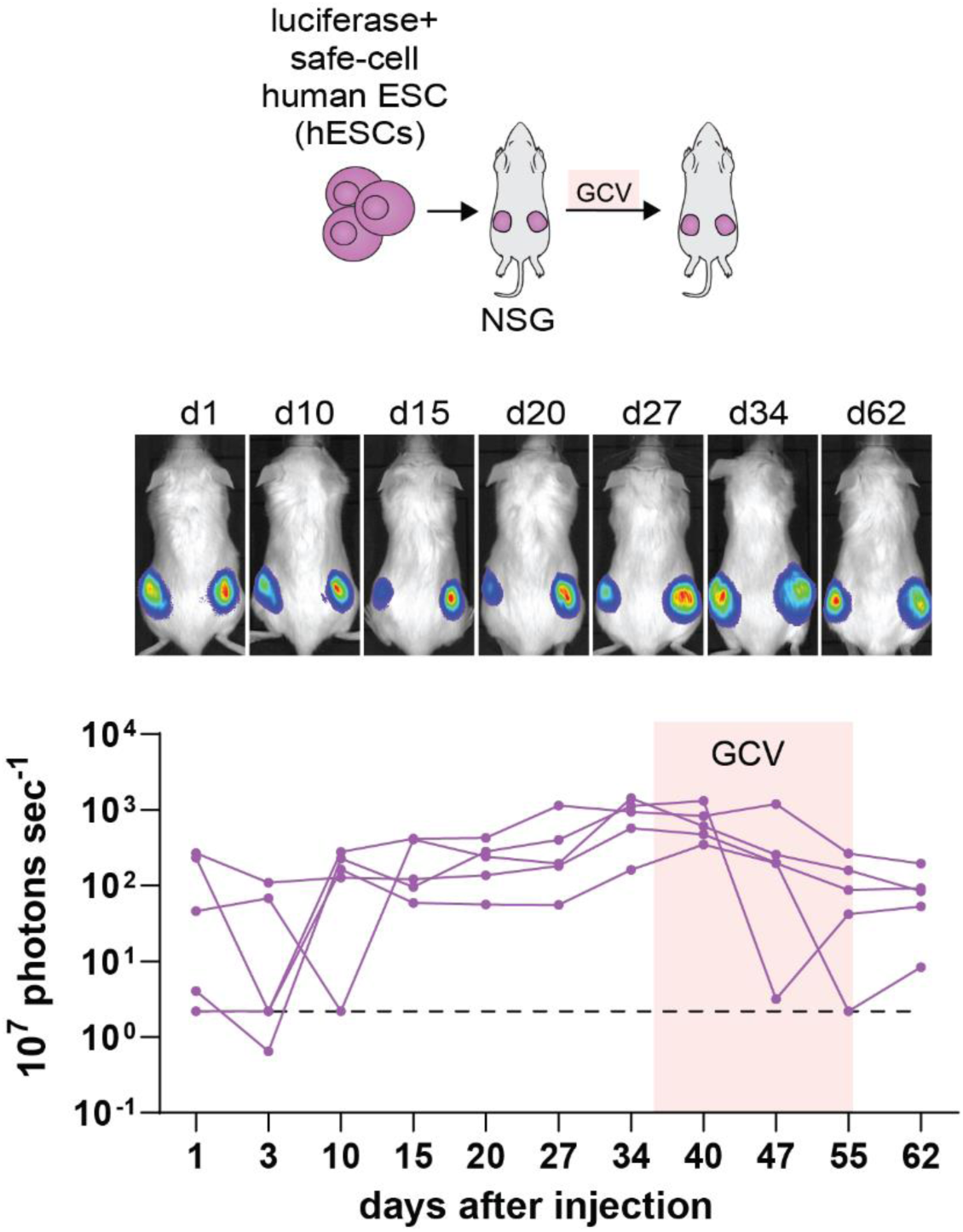
Safe-cell hESCs-derived teratomas in NSG mice. 10-12 million luciferase+ safe-cell human ESCs were injected subcutaneously into each flanks of NSG mice and tracked for 62 days by BLI. Recipients were treated with GCV to halt safe-cell hESC growth. Images show one representative mouse, graph shows BLI photon emission of all mice. Black dotted line in graph shows detection limit. BLI exposure time; 10 seconds.

## References

1. Taylor, C.J., Peacock, S., Chaudhry, A.N., Bradley, J.A. & Bolton, E.M. Generating an iPSC bank for HLA-matched tissue transplantation based on known donor and recipient HLA types. Cell Stem Cell 11, 147–152 (2012).

2. Gornalusse, G.G. et al. HLA-E-expressing pluripotent stem cells escape allogeneic responses and lysis by NK cells. Nat Biotechnol 35, 765–772 (2017).

3. Deuse, T. et al. Hypoimmunogenic derivatives of induced pluripotent stem cells evade immune rejection in fully immunocompetent allogeneic recipients. Nat Biotechnol 37, 252–258 (2019).

4. Han, X. et al. Generation of hypoimmunogenic human pluripotent stem cells. Proc Natl Acad Sci U S A 116, 10441–10446 (2019).

5. Warren, E.H. et al. Effect of MHC and non-MHC donor/recipient genetic disparity on the outcome of allogeneic HCT. Blood 120, 2796–2806 (2012).

6. Spierings, E. Minor histocompatibility antigens: past, present, and future. Tissue Antigens 84, 374–360 (2014).

7. Zhang, Q. & Reed, E.F. The importance of non-HLA antibodies in transplantation. Nat Rev Nephrol 12, 484–495 (2016).

8. Ostrander, E.A., Davis, B.W. & Ostrander, G.K. Transmissible Tumors: Breaking the Cancer Paradigm. Trends Genet 32, 1–15 (2016).

9. Pye, R.J. et al. A second transmissible cancer in Tasmanian devils. Proc Natl Acad Sci U S A 113, 374–379 (2016).

10. Stammnitz, M.R. et al. The Origins and Vulnerabilities of Two Transmissible Cancers in Tasmanian Devils. Cancer Cell 33, 607–619 e615 (2018).

11. Liang, Q. et al. Linking a cell-division gene and a suicide gene to define and improve cell therapy safety. Nature 563, 701–704 (2018).

12. Forster, R., Davalos-Misslitz, A.C. & Rot, A. CCR7 and its ligands: balancing immunity and tolerance. Nat Rev Immunol 8, 362–371 (2008).

13. Shields, J.D., Kourtis, I.C., Tomei, A.A., Roberts, J.M. & Swartz, M.A. Induction of lymphoidlike stroma and immune escape by tumors that express the chemokine CCL21. Science 328, 749–752 (2010).

14. Sharpe, A.H. & Pauken, K.E. The diverse functions of the PD1 inhibitory pathway. Nat Rev Immunol 18, 153–167 (2018).

15. Bouillet, P. & O’Reilly, L.A. CD95, BIM and T cell homeostasis. Nat Rev Immunol 9, 514–519 (2009).

16. Smyth, M.J., Sullivan, L.C., Brooks, A.G. & Andrews, D.M. Non-classical MHC Class I molecules regulating natural killer cell function. Oncoimmunology 2, e23336 (2013).

17. Sun, J. et al. A cytosolic granzyme B inhibitor related to the viral apoptotic regulator cytokine response modifier A is present in cytotoxic lymphocytes. J Biol Chem 271, 27802–27809 (1996).

18. Matozaki, T., Murata, Y., Okazawa, H. & Ohnishi, H. Functions and molecular mechanisms of the CD47-SIRPalpha signalling pathway. Trends Cell Biol 19, 72–80 (2009).

19. Nathan, C. & Muller, W.A. Putting the brakes on innate immunity: a regulatory role for CD200? Nat Immunol 2, 17–19 (2001).

20. Brissette, M.J. et al. MFG-E8 released by apoptotic endothelial cells triggers anti-inflammatory macrophage reprogramming. PLoS One 7, e36368 (2012).

21. Ding, S. et al. Efficient transposition of the piggyBac (PB) transposon in mammalian cells and mice. Cell 122, 473–483 (2005).

22. Cadinanos, J. & Bradley, A. Generation of an inducible and optimized piggyBac transposon system. Nucleic Acids Res 35, e87 (2007).

23. Zhao, S. et al. PiggyBac transposon vectors: the tools of the human gene encoding. Transl Lung Cancer Res 5, 120–125 (2016).

24. Ivics, Z., Hackett, P.B., Plasterk, R.H. & Izsvak, Z. Molecular reconstruction of Sleeping Beauty, a Tc1-like transposon from fish, and its transposition in human cells. Cell 91, 501–510 (1997).

25. Izsvak, Z. & Ivics, Z. Sleeping beauty transposition: biology and applications for molecular therapy. Mol Ther 9, 147–156 (2004).

26. Niwa, H., Yamamura, K. & Miyazaki, J. Efficient selection for high-expression transfectants with a novel eukaryotic vector. Gene 108, 193–199 (1991).

27. Gertsenstein, M. et al. Efficient generation of germ line transmitting chimeras from C57BL/6N ES cells by aggregation with outbred host embryos. PLoS One 5, e11260 (2010).

28. Tan, Y. et al. MFG-E8 Is Critical for Embryonic Stem Cell-Mediated T Cell Immunomodulation. Stem Cell Reports 5, 741–752 (2015).

29. Abdullah, Z. et al. Serpin-6 expression protects embryonic stem cells from lysis by antigen-specific CTL. J Immunol 178, 3390–3399 (2007).

30. Horn, C. et al. Splinkerette PCR for more efficient characterization of gene trap events. Nat Genet 39, 933–934 (2007).

31. Woltjen, K. et al. piggyBac transposition reprograms fibroblasts to induced pluripotent stem cells. Nature 458, 766–770 (2009).

32. Rong, Z. et al. An effective approach to prevent immune rejection of human ESC-derived allografts. Cell Stem Cell 14, 121–130 (2014).

33. Lanza, R., Russell, D.W., and Nagy, A Engineering universal cells that evade immune detection. Nature Reviews Immunology In Press (2019).

34. Falkenburg, J.H., van de Corput, L., Marijt, E.W. & Willemze, R. Minor histocompatibility antigens in human stem cell transplantation. Exp Hematol 31, 743–751 (2003).

35. King, A. et al. Surface expression of HLA-C antigen by human extravillous trophoblast. Placenta 21, 376–387 (2000).

36. Bolger, A.M., Lohse, M. & Usadel, B. Trimmomatic: a flexible trimmer for Illumina sequence data. Bioinformatics 30, 2114–2120 (2014).

37. Bray, N.L., Pimentel, H., Melsted, P. & Pachter, L. Near-optimal probabilistic RNA-seq quantification. Nat Biotechnol 34, 525–527 (2016).

38. Love, M.I., Huber, W. & Anders, S. Moderated estimation of fold change and dispersion for RNA-seq data with DESeq2. Genome Biol 15, 550 (2014).

39. Dobin, A. et al. STAR: ultrafast universal RNA-seq aligner. Bioinformatics 29, 15–21 (2013).

40. Robinson, J.T. et al. Integrative genomics viewer. Nat Biotechnol 29, 24–26 (2011).

41. Thorvaldsdottir, H., Robinson, J.T. & Mesirov, J.P. Integrative Genomics Viewer (IGV): high-performance genomics data visualization and exploration. Brief Bioinform 14, 178–192 (2013).

