## Supplementary Tables for "Induction of long-term allogeneic cell acceptance and formation of immune privileged tissue in immunocompetent hosts"

| donor mESC --> recipient strain | cage ID | # injected recipients | # recipients that develop teratomas | length of observation |
|---|---|---|---|---|
| **safe-cell C57BL/6 mESCs --> C57BL/6** | C7003871 | 2 | 1 | 2 months |
| | C7003902 | 2 | 2 | 2 months |
| | C7003790 | 5 | 5 | 3 months |
| | C7170476 | 5 | 5 | 3 months |
| | C7212976 | 5 | 4 | 3 months |
| | C7283968 | 3 | 3 | 3 months |
| | C7658219 | 5 | 5 | 6 months |
| | C7658226 | 5 | 4 | 6 months |
| | C7720221 | 5 | 5 | 2 months |
| | C7658242 | 5 | 5 | 2 months |
| | C7912243 | 3 | 3 | 6 months |
| | C7912258 | 3 | 3 | 6 months |
| | C8200160 | 5 | 2 | 4 months |
| | C8200187 | 4 | 3 | 4 months |
| | C8261223 | 5 | 3 | 4 months |
| | C8261206 | 5 | 5 | ongoing |
| | C8321799 | 5 | 3 | ongoing |
| | C8321813 | 5 | 4 | ongoing |
| **Klg-1 safe-cell (C57BL/6) mESCs --> C57BL/6** | C7003800 | 5 | 5 | 5 months |
| | C7003788 | 5 | 5 | 5 months |
| | C7003939 | 3 | 3 | 2 months |
| | C7003863 | 5 | 5 | 6 months |
| | C7212969 | 5 | 5 | 5 months |
| | C7212982 | 5 | 5 | 6 months |
| | C7283975 | 5 | 5 | 4 months |
| | C7283981 | 5 | 5 | 4 months |
| | C7283952 | 5 | 5 | 5 months |
| | C7420625 | 3 | 3 | 5 months |
| | C7524032 | 5 | 5 | 6 months |
| | C7524050 | 5 | 5 | 6 months |
| | C7720213 | 5 | 5 | 2 months |
| | C7658235 | 5 | 5 | 2 months |
| | C7749067 | 5 | 5 | 2 months |
| | C8200194 | 4 | 4 | ongoing |
| | C8185883 | 5 | 4 | 4 months |

| | | | | |
|---|---|---|---|---|
| | C8261186 | 5 | 5 | ongoing |
| | C8261247 | 5 | 4 | ongoing |
| | C8321752 | 5 | 5 | ongoing |
| | C8321809 | 5 | 5 | ongoing |
| **Klg-1 safe-cell (C57BL/6) mESCs--> FVB** | C7003676 | 5 | 5 | 2 months |
| | C7003703 | 5 | 5 | 9 months |
| | C7003742 | 4 | 3 | 4 months |
| | C7143911 | 4 | 4 | 7 months |
| | C7238930 | 4 | 4 | 6 months |
| | C7308893 | 2 | 1 | 2 months |
| | C7308903 | 3 | 0 | n/a |
| | C7698704 | 5 | 5 | 5 months |
| | C7924439 | 5 | 4 | 3 months |
| | C7924456 | 5 | 2 | 4 months |
| | C7667371 | 3 | 3 | 6 months |
| | C7667385 | 5 | 3 | 7 months |
| | C7766817 | 3 | 2 | 4 months |
| | C8268940 | 5 | 2 | 3 months |
| | C8268938 | 5 | 4 | ongoing |
| | C8334503 | 5 | 1 | ongoing |
| | C8334542 | 5 | 5 | ongoing |
| **Klg-1 safe-cell (C57BL/6) mESCs --> C3H** | C7434570 | 5 | 3 | 4 months |
| | C7744419 | 3 | 3 | 2 months |
| | C7685455 | 4 | 2 | 4 months |
| | C7744403 | 3 | 2 | 4 months |
| | C7685440 | 2 | 2 | 5 months |
| | C7633765 | 5 | 1 | 4 months |
| | C7633754 | 4 | 4 | 4 months |
| | C8269025 | 4 | 3 | 3 months |
| | C8269039 | 5 | 5 | 3 months |
| | C8213761 | 5 | 3 | 3 months |
| | C8334463 | 5 | 5 | 3 months |
| | C8334471 | 5 | 5 | ongoing |
| **Klg-1 safe-cell (C57BL/6) mESCs --> BALB/c** | C8542655 | 5 | 4 | 4 months |
| | C8542638 | 5 | 4 | 4 months |
| | C8542640 | 5 | 0 | no teratomas |
| | C8542629 | 5 | 4 | 4 months |
| **Klg-1 safe-cell (C57BL/6) mESCs --> CD1** | C7170495 | 5 | 3 | 7 months |
| | C7170482 | 5 | 0 | n/a |
| | C7283934 | 4 | 3 | 5 months |
| | C7283947 | 5 | 1 | 6 months |
| | C7215283 | 5 | 2 | 8 months |
| | C7283923 | 3 | 0 | no teratomas |
| | C7436427 | 5 | 0 | no teratomas |

| | | | | |
|---|---|---|---|---|
| **Klg-2 safe-cell (C57BL/6) mESCs --> C57BL/6** | C8185877 | 5 | 5 | ongoing |
| | C8185865 | 5 | 5 | ongoing |
| | C7524045 | 5 | 5 | 6 months |
| | C8261210 | 5 | 5 | 3 months |
| | C8261193 | 5 | 5 | ongoing |
| | C8321781 | 5 | 3 | ongoing |
| | C8321775 | 5 | 3 | 3 months |
| **Klg-2 safe-cell (C57BL/6) mESCs --> FVB** | C7459341 | 2 | 1 | 3 months |
| | C7496163 | 5 | 3 | 6 months |
| | C7443801 | 5 | 0 | no teratomas |
| | C8116956 | 5 | 0 | no teratomas |
| | C8268964 | 5 | 0 | no teratomas |
| | C8268972 | 5 | 0 | no teratomas |
| | C8334492 | 5 | 1 | 3 months |
| | C8334519 | 5 | 0 | no teratomas |
| **Klg-2 safe-cell (C57BL/6) mESCs --> C3H** | C7538143 | 5 | 3 | 4 months |
| | C7538136 | 5 | 3 | 4 months |
| | C7434558 | 5 | 2 | 5 months |
| | C8135726 | 2 | 2 | 6 months |
| | C7967151 | 2 | 2 | 4 months |
| | C7848218 | 2 | 2 | 6 months |
| | C8213816 | 5 | 2 | 5 months |
| | C8213828 | 5 | 2 | 5 months |
| | C8213837 | 5 | 4 | 5 months |
| | C8268986 | 4 | 2 | 4 months |
| | C8269002 | 5 | 0 | no teratomas |
| | C8334485 | 5 | 4 | 3 months |
| | C8334437 | 5 | 1 | ongoing |

**Supplementary Table S1 |** Tabulation of all experiments in which Klg-1 and Klg-2 mESCs were transplanted into isogeneic and allogeneic recipients and monitored teratoma formation and persistence. Table shows number of recipients injected, number of recipients that formed teratomas, and the length of observation before euthanasia for each cage among all transplantation. Data corresponds to Fig. 3c-e in main text for safe-cell and Klg-1 mESCs, and Supplementary Fig. S3 for Klg-2 mESCs.

| | chr# | start | end | associated gene (basal + extension) | associated gene (1kb upstream, 10kb downstream) | class |
|---|---|---|---|---|---|---|
| 1 | chr1 | 191717633 | 191717637 | Lpgat1 (-754) | Lpgat1 (-754) | a |
| 2 | chr1 | 134405686 | 134405690 | Cyb5r1 (-93) | Cyb5r1 (-93) | a |
| 3 | chr1 | 84829501 | 84829505 | Dner (-133282), Trip12 (+9801) | Trip12 (+9801) | a |
| 4 | chr1 | 23153142 | 23153146 | 4933415F23Rik (-50891), Gm6420 (+103262) | NONE | a |
| 5 | chr2 | 180071074 | 180071078 | Gtpbp5 (-717) | Gtpbp5 (-717) | a |
| 6 | chr5 | 107123013 | 107123017 | Cdc7 (+158454), Tgfbr3 (+166614) | NONE | a |
| 7 | chr5 | 73011177 | 73011181 | Slc10a4 (+4275), Fryl (+245409) | Slc10a4 (+4275) | a |
| 8 | chr6 | 32083096 | 32083100 | 1700012A03Rik (+32852), Plxna4 (+505094) | NONE | a |
| 9 | chr6 | 16840125 | 16840129 | Tfec (+58314) | NONE | a |
| 10 | chr7 | 138527843 | 138527847 | Tcerg1l (-130117), Mapk1ip1 (+318428) | NONE | a |
| 11 | chr7 | 114902623 | 114902627 | Gm6816 (+25101), Insc (+156859) | NONE | a |
| 12 | chr7 | 36768266 | 36768270 | Tshz3 (+70150) | NONE | a |
| 13 | chr8 | 124648197 | 124648201 | 2310022B05Rik (+15170), Capn9 (+72088) | NONE | a |
| 14 | chr10 | 113185466 | 113185470 | Atxn7l3b (-256442) | NONE | a |
| 15 | chr10 | 67187286 | 67187290 | Jmjd1c (+60030), Nrbf2 (+98017) | NONE | a |
| 16 | chr11 | 72859864 | 72859868 | Atp2a3 (-101303), Zzef1 (+63612) | NONE | a |
| 17 | chr12 | 72744695 | 72744699 | Dhrs7 (-79869), Ppm1a (-16514) | NONE | a |
| 18 | chr14 | 76625222 | 76625226 | Serp2 (-68537), Lacc1 (+411417) | NONE | a |
| 19 | chr14 | 70749859 | 70749863 | Npm2 (-96777), Xpo7 (+16767) | NONE | a |
| 20 | chr15 | 74654046 | 74654050 | Mroh4 (-17787), Arc (+18520) | NONE | a |
| 21 | chr16 | 20360201 | 20360205 | Cyp2ab1 (-37138), Abcc5 (+66191) | NONE | a |
| 22 | chr17 | 33355492 | 33355496 | Zfp81 (+3384), Zfp955b (+65845) | Zfp81 (+3384) | a |
| 23 | chr18 | 23803174 | 23803178 | Mapre2 (-808) | Mapre2 (-808) | a |
| 24 | chr19 | 55896945 | 55896949 | Habp2 (-390975), Tcf7l2 (+155137) | NONE | a |
| 25 | chr19 | 5327933 | 5327937 | Catsper1 (-7806), Gal3st3 (+29604) | NONE | a |
| 26 | chr1 | 161781388 | 161781392 | Fasl (+7105), Tnfsf18 (+286735) | Fasl (+7105) | c |
| 27 | chr1 | 132237685 | 132237689 | Klhdc8a (-60939), Gm10188 (-7920) | NONE | c |
| 28 | chr2 | 50914093 | 50914097 | Mmadhc (-617378), Rnd3 (+235016) | NONE | c |
| 29 | chr3 | 98013125 | 98013129 | Notch2 (-411) | Notch2 (-411) | c |
| 30 | chr4 | 41129612 | 41129616 | Ube2r2 (-6129), Nol6 (-5159) | NONE | c |
| 31 | chr8 | 117017967 | 117017971 | Gcsh (-24432), Pkd1l2 (+64480) | NONE | c |
| 32 | chr10 | 38610142 | 38610146 | Rfpl4b (+211635) | NONE | c |
| 33 | chr11 | 120598854 | 120598858 | Alyref (-491), Anapc11 (+435) | Alyref (-491), Anapc11(+435) | c |
| 34 | chr11 | 88966965 | 88966969 | Scpep1 (-11502), Coil (-6968) | NONE | c |

| | | | | | | |
|---|---|---|---|---|---|---|
| 35 | chr11 | 31392342 | 31392346 | Stc2 (-22270), Bod1 (+279541) | NONE | c |
| 36 | chr11 | 15569072 | 15569076 | Vstm2a (-688650) | NONE | c |
| 37 | chr12 | 82470236 | 82470240 | Gm5435 (+26299), Sipa1l1 (+158889) | NONE | c |
| 38 | chr12 | 69177378 | 69177382 | Rpl36al (+6633), Lrr1 (+8566) | Rpl36al (+6633), Lrr1 (+8566) | c |
| 39 | chr18 | 40060860 | 40060864 | Yipf5 (+158537), Pabpc2 (+287365) | NONE | c |
| 40 | chrX | 87579696 | 87579700 | Il1rapl1 (+535947) | NONE | c |
| 41 | chr1 | 128441620 | 128441624 | Dars (-24206), Cxcr4 (+150668) | NONE | b |
| 42 | chr1 | 86237174 | 86237178 | B3gnt7 (-66045), Armc9 (+82366) | NONE | b |
| 43 | chr1 | 47274114 | 47274118 | Slc39a10 (-420070) | NONE | b |
| 44 | chr2 | 129419766 | 129419770 | Il1b (-48629), F830045P16Rik (+116834) | NONE | b |
| 45 | chr2 | 129178651 | 129178655 | Slc20a1 (-20111), Chchd5 (+48953) | NONE | b |
| 46 | chr2 | 17067679 | 17067683 | Nebl (+663362), Plxdc2 (+711377) | NONE | b |
| 47 | chr3 | 64374755 | 64374759 | Vmn2r3 (-87340), Vmn2r4 (+35300) | NONE | b |
| 48 | chr3 | 30939259 | 30939263 | Phc3 (+30140), Gpr160 (+83311) | NONE | b |
| 49 | chr5 | 115001292 | 115001296 | Hnf1a (-30227), Sppl3 (-9843) | NONE | b |
| 50 | chr7 | 92870412 | 92870416 | Prcp (-4839), 4632434I11Rik (+3799) | 4632434I11Rik (+3799) | b |
| 51 | chr13 | 45691011 | 45691015 | Gmpr (+183569), Atxn1 (+273978) | NONE | b |
| 52 | chr14 | 14308741 | 14308745 | Olfr720 (-132626), Olfr31 (-19370) | NONE | b |
| 53 | chr15 | 35305533 | 35305537 | Vps13b (-66011), Osr2 (+9437) | Osr2 (+9437) | b |
| 54 | chr18 | 67161288 | 67161292 | Gnal (+72954), Mppe1 (+84540) | NONE | b |
| 55 | chr19 | 38967824 | 38967828 | Cyp2c55 (-39193), Hells (+36911) | NONE | b |
| 56 | chrX | 65503194 | 65503198 | Ctag2 (+455552), 4930447F04Rik (+801028) | NONE | b |

**Supplementary Table S2 |** Number and position of *piggyBac* insertions, each containing an immunomodulatory transgene, in Klg-1 mESCs. Data was generated using splinkerette PCR with primers against the long terminal repeats of *piggyBac* flanking and comparing the next generation (NGS)-sequenced amplified products to a mouse reference genome. Complete description in methods. Table classes defined by number of reads on each TTAA-flanking side. All classes have greater than 10 reads on one side; class a, one read on one other side; class b, 2-4 reads on other one side; class c, more than 4 reads on other side one side.

| Gene | NCIB gene ID | Forward Primer for gateway cloning (attb1 site in black) | Reverse primer for gateway cloning (attb2 site in black) |
|---|---|---|---|
| *Pdl1* | 60533 | GGGACAAGTTTGTACAAAAAAGCAGCCT ATGAGGATATTTGCTGGCATTATAT | GGGACCACTTTGTACAAGAAAGCTGGGT TTACGTCTCCTCGAATTGTGTATCA |
| *Cd200* | 17470 | GGGACAAGTTTGTACAAAAAAGCAGCCT TAATGAAAGTCGACGGCGCGCCGAA | GGGACCACTTTGTACAAGAAAGCTGGGT AATTAACCCTCACTAAAGGGACCTA |
| *Cd47* | 16423 | GGGACAAGTTTGTACAAAAAAGCAGCCT GTGACACTATAGAACAAGTT | GGGACCACTTTGTACAAGAAAGCTGGGT AGTACAAGAAAGCTGGGTACG |
| *H2-M3* | 14991 | GGGACAAGTTTGTACAAAAAAGCAGCCT CGTCGACCCACGCGTCCGCAGGGC | GGGACCACTTTGTACAAGAAAGCTGGGT GAGGGATACTCTAGAGCGGCCGCTT |
| *Fasl* | 14103 | GGGACAAGTTTGTACAAAAAAGCAGCCT CCTGCAGGTACCGGTCCGGAATTCC | GGGACCACTTTGTACAAGAAAGCTGGGT CACGCGTACGTAAGCTTGGATCCTC |
| *Mfge8* | 17304 | GGGACAAGTTTGTACAAAAAAGCAGCCT GTACAAAAAAAGCAGGCTGGTACC | GGGACCACTTTGTACAAGAAAGCTGGGT ACAGCCCAGCAGCTCCAGGCGCAGGGT |
| *Serpinb9* | 20723 | n/a | n/a |
| *Ccl21* | 18829 | n/a | n/a |

**Supplementary Table S3 |** Attb1-forward and attb2-reverse primers used for gateway cloning.

| Gene | Primer |
|---|---|
| Pdl1 | ATCATCCCAGAACTGCCTGC |
| Cd200 | TCTCTCTCTCTCTCCCTCCC |
| Cd47 | CAGTGCACCCCAGTTTTGAA |
| H2-M3 | GGGGATCCAGGACTTCTTCA |
| Fasl | AGAGGGATGTTTGAGGCACA |
| Mfge8 | TTCTGTGACTCCAGCCTGTG |
| Serpinb9 | CACCTTTGCCATCCATCTTT |
| Ccl21a | TCCGAGGCTATAGGAAGCAA |

**Supplementary Table S4 |** Primers used to verify transgene sequence after cloning into gateway pDONr221 vectors.

| Gene | Forward Primer | Reverse Primer |
|---|---|---|
| *Pdl1* | GCGAATCACGCTGAAAGTCAATGC | ACGGGTTGGTGGTCACTGTTTG |
| *Cd200* | CTCTCCACCTACAGCCTGATT | AGAACATCGTAAGGATGCAGTTG |
| *Cd47* | TGCGGTTCAGCTCAACTACTG | GCTTTGCGCCTCCACATTAC |
| *H2-M3* | ACTATCAGGCTGCGTATGATGG | ACTCTAAACGGCTCTTCGTGA |
| *Fasl* | TGGTTGGAATGGGATTAGGA | GTGGGGGTTCCCTGTTAAAT |
| *Mfge8* | CCTGGGCCTGAAGAATAACA | GCATTGATCTTGCCCTGATT |
| *Serpinb9* | TAGAGTGGCCAACAGGCTCT | GGAGCCACCTGACAACAACT |
| *Ccl21* | GTGATGGAGGGGGTCAGGA | GGGATGGGACAGCCTAAACT |
| *Eef2* | GTATTAAGAGCTGCGACCCC | ATAGAAGCGGCCTTTGTCAG |

**Supplementary Table S5 |** Forward and reverse primers used to detect immunomodulatory transgenes, as well as a control gene *Eef2* in mESCs clones by RT-qPCR.
